# DNMT1 overexpression disrupts DNA methylation homeostasis in mouse embryonic stem cells

**DOI:** 10.64898/2026.01.26.701550

**Authors:** Elizabeth Elder, Anthony Lemieux, Serge McGraw

## Abstract

DNMT1 overexpression is frequently observed in cancer and other pathologies, yet its direct impact on DNA methylation remains poorly defined. Using whole-genome methyl sequencing, we show that overexpressing DNMT1 in mouse embryonic stem cells (mESCs) induces global hypomethylation, focal hypermethylation in promoters and CpG islands, and increased methylome variability—well-documented characteristics of cancer. We also find that differential promoter methylation is associated with developmental (axon guidance, Wnt) and disease (cancer, cardiomyopathy) pathways and correlates with altered gene expression. Additionally, DNMT3A/B levels are reduced, indicating that excess DNMT1 perturbs the broader DNA methylation machinery. Moreover, to model targeted therapies, we reveal that hypermethylation is mostly erased following DNMT1 depletion, but a substantial portion persists. Hypermethylation is then largely regained upon reinstating DNMT1 overexpression, only achieving permanent erasure at a minority of regions. Finally, promoter hypermethylation detected in mESCs is observed across diverse human cancers, supporting its biological significance and the translational relevance of this study. Overall, these findings illuminate how DNMT1 overexpression disrupts DNA methylation homeostasis, providing mechanistic insight into its pathogenic consequences and therapeutic targeting.

**Graphical abstract:** 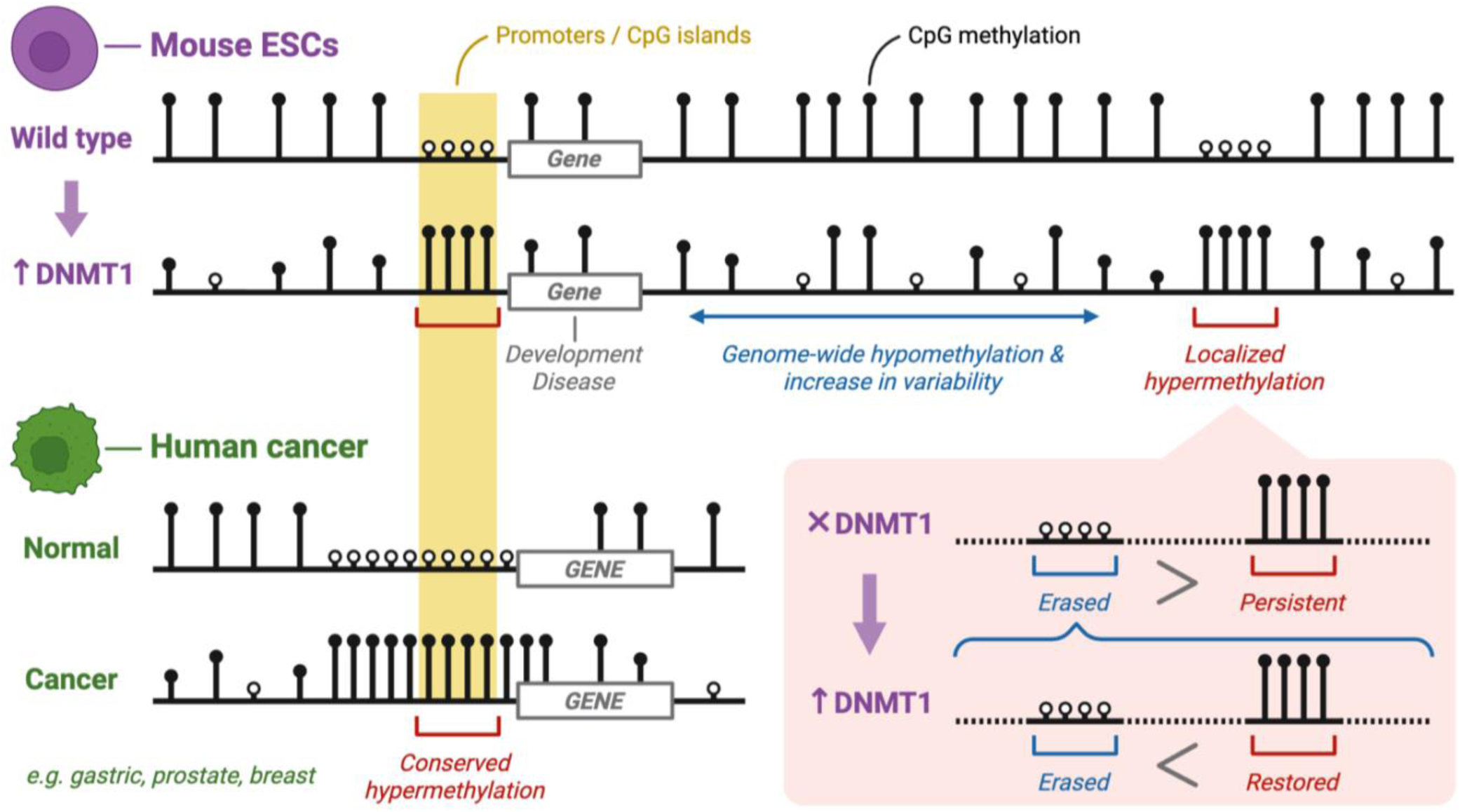

## Introduction

DNA methylation is one of the most studied epigenetic mechanisms in development and disease, regulating gene expression, chromatin structure, genome stability, X-chromosome inactivation, genomic imprinting and transposable element silencing^1^. In mammals, DNA methylation occurs predominantly at cytosines within CpG (cytosine-phosphate-guanine) sites and is regulated by DNA methyltransferase (DNMT) enzymes. DNMT3A and DNMT3B mediate de novo methylation and DNMT1 maintains methylation patterns during DNA replication^1^, though evidence supports that functional overlap can occur where DNMTs act outside their canonical roles. It has been reported that, in some contexts, DNMT1 can contribute to de novo methylation^2–4^, and DNMT3A and DNMT3B to maintenance^5–7^. The coordinated activity of these enzymes ensures faithful propagation of DNA methylation patterns while permitting dynamic remodeling during cell differentiation^8^ and in response to physiological^9,10^ and environmental cues^11,12^. DNA methylation regulation depends not only on the catalytic activities of individual DNMTs but also on their interaction with other epigenetic regulators^13–16^ and, importantly, on their relative abundance. DNMT dosage fluctuates in a tightly controlled manner throughout development^17^ and the cell cycle^18^ in order to precisely tune DNA methylation patterns to the evolving regulatory programs of proliferating, differentiating and lineage-committed cells.

Abnormal DNMT activity, whether through deficiency or excess, can lead to aberrant DNA methylation patterns and has been linked to various pathologies, including developmental syndromes^19,20^, neurological diseases^21,22^, imprinting disorders^23^ and cancer^24^. For example, in cancer, where DNMTs are often overexpressed^24^, global hypomethylation, focal hypermethylation in promoters and CpG islands, and increased CpG-to-CpG variability are recurrent signatures^25,26^. Because disease-associated DNA methylation alterations show sufficient stability and specificity, they are widely studied as biomarkers for diagnosis, prognosis and treatment stratification^27^; some DNA methylation-based tests are already in clinical use, such as the SEPT9 gene methylation assay for colorectal cancer screening^28^. Moreover, DNMTs have become attractive therapeutic targets, particularly in cancer to counteract their overexpression. Pan-DNMT inhibitors (e.g. azacitidine, decitabine) are approved for the treatment of hematological malignancies, and ongoing efforts aim to extend their application to solid tumors^29^. More recently, DNMT1-selective inhibitors (e.g. GSK3685032, GSK3482364) have emerged as promising alternatives that may offer enhanced efficacy with fewer systemic side effects than pan-inhibiting agents^30^. Despite these advances, specific mechanisms through which DNMT overexpression contributes to cancer methylomes remain elusive, as they can be obscured by the extensive intratumoral and interpatient heterogeneity and by the multitude of concurrent oncogenic processes. Paradoxical mechanisms especially, such as DNMT overexpression coexisting with global hypomethylation in cancer, are difficult to resolve in highly complex biological systems.

Overexpression of DNMT1 can be studied in a more direct and controlled setting thanks to the transgenic *Dnmt1^tet/tet^* mouse embryonic stem cells (mESCs), in which *Dnmt1* expression is regulated by the Tet-Off system^31^. The Tet-Off system is typically used to reversibly repress a gene through doxycycline treatment and recovery; for instance, we have previously employed *Dnmt1^tet/tet^* mESCs to investigate the reversibility of DNMT1 loss^32,33^. However, due to the strong promoter-transactivator pairing in the Tet-Off construct, untreated *Dnmt1^tet/tet^* mESCs robustly exhibit elevated DNMT1 levels compared to wild-type mESCs^31^. *Dnmt1^tet/tet^* mESCs have therefore been used to study DNMT1 overexpression, but only in developmental and neurological disease contexts thus far^31,34–36^, leaving their potential to illuminate cancer-related mechanisms unexplored. While prior studies using *Dnmt1^tet/tet^* or other DNMT1-overexpressing mESCs have associated DNMT1 overexpression almost exclusively with DNA hypermethylation or reported no measurable effect on DNA methylation, their conclusions were drawn from locus-specific DNA methylation analyses^31,37^ or from incomplete or low-resolution views of the methylome generated by RRBS (reduced-representation bisulfite sequencing)^36^, reverse-phase HPLC (high-performance liquid chromatography)^37^ or methylation-sensitive enzymatic restriction coupled with Southern blotting^37^. Fully characterizing how DNMT1 overexpression reshapes the mESC methylome would require a whole-genome, high-resolution detection method such as enzymatic methyl sequencing (EM-seq)^38^ performed with adequate sequencing depth (i.e. ≥ 10X per CpG).

Here, using high-depth EM-seq, we comprehensively profile DNA methylation in *Dnmt1^tet/tet^* versus wild-type mESCs, uncovering compelling parallels with cancer methylomes, including genome-wide hypomethylation, localized hypermethylation and higher CpG-to-CpG variability. We show that DNA methylation alterations in promoters are consistent with the diverse pathological potential of DNMT1 overexpression and correlate with changes in gene expression. Moreover, by examining how excess DNMT1 affects other components of the DNA methylation machinery and integrating binding sites of transcriptional regulators, we propose candidate underlying mechanisms that warrant further investigation. We also assess the impact of acutely depleting DNMT1 in *Dnmt1^tet/tet^* mESCs and subsequently reinstating its overexpression, providing insight relevant to DNMT-inhibitor therapies. Finally, comparative analyses with human cancer DNA methylation datasets show the translational relevance of our findings.

## Results and Discussion

### Perturbation of the DNA methylation machinery in *Dnmt1^tet/tet^* mESCs

To investigate how excess DNMT1 influences DNA methylation regulation, we employed the transgenic *Dnmt1^tet/tet^* mESCs developed by Borowczyk et al., which overexpress DNMT1^31^ (Figure 1A). *Dnmt1* gene expression was increased 7-fold relative to wild-type mESCs, while pluripotency (*Pou5f1*, *Sox2*, *Nanog*) and housekeeping (*Actb*, *Gapdh*, *Sdha*) genes remained unchanged (Figure 1B; Data S1). At the protein level, DNMT1 was elevated 3-fold, with ACTB (beta-actin) levels unaffected (Figures 1C–D, S1; Data S2). In addition to stable expression of pluripotency genes, *Dnmt1^tet/tet^* mESCs exhibited unlimited proliferation and robust colony formation, indicating overall preservation of stemness as previously reported^33–36^. We profiled methylation at CpG sites using EM-seq, revealing a global loss of 8% in *Dnmt1^tet/tet^* versus wild-type mESCs: mean levels were 76% in wild-type and 68% in *Dnmt1^tet/tet^* mESCs, respectively (Figure 1E).

**Figure 1.**
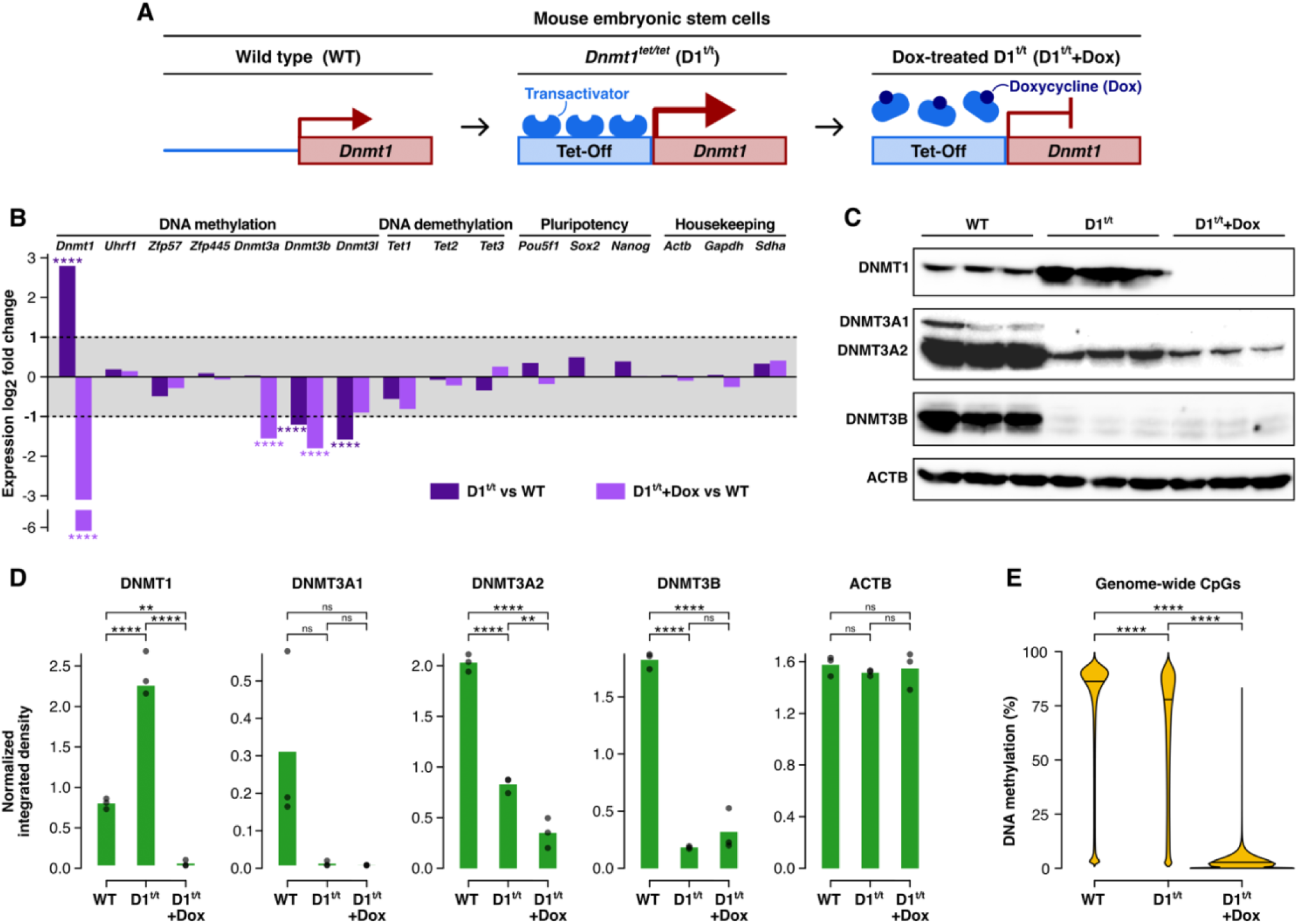
Perturbation of the DNA methylation machinery in *Dnmt1^tet/tet^* mESCs. **(A)** The *Dnmt1^tet/tet^* mouse embryonic stem cell (mESC) model in which *Dnmt1* expression is controlled by the Tet-Off system. **(B)** Gene expression log_2_ fold change in *Dnmt1^tet/tet^* and doxycycline-treated *Dnmt1^tet/tet^* versus wild-type mESCs. Gene expression was measured by mRNA-seq (n=3 per condition) and quantified using the DESeq2 R package. Statistical significance was defined by a log_2_ fold change ≤ -1 or ≥ 1 and an adjusted p-value ≤ 0.05. *See also Data S1*. **(C)** Protein levels measured by western blot in wild-type, *Dnmt1^tet/tet^* and doxycycline-treated *Dnmt1^tet/tet^* mESCs (n=3 per condition). **(D)** Quantification of western blot bands in **C** using ImageJ software (normalized to Ponceau S staining). Statistical comparisons were conducted using one-way ANOVA with Tukey’s HSD test and p-value adjustment. *See also Figure S1 and Data S2*. **(E)** DNA methylation levels of genome-wide CpGs in wild-type, *Dnmt1^tet/tet^* and doxycycline-treated *Dnmt1^tet/tet^* mESCs measured by EM-seq (n=3 per condition). Statistical comparisons were conducted using the pairwise Wilcoxon test with Benjamini-Hochberg p-value adjustment. ** p-value ≤ 0.01, **** p-value ≤ 0.0001, ns: not significant.

Global hypomethylation in *Dnmt1^tet/tet^* mESCs was unexpected as it opposes prior studies that associated DNMT1 overexpression in mESCs with hypermethylation or with no changes at all. However, these studies either emphasized locus-specific alterations^31,37^ or relied on detection assays (i.e. RRBS^36^, reverse-phase HPLC^37^, methylation-sensitive enzymatic restriction coupled with Southern blotting^37^) that have important limitations compared to EM-seq regarding genomic coverage, sensitivity, resolution, or CpG density bias. Hypomethylation in *Dnmt1^tet/tet^* mESCs may have resulted from excess DNMT1 promoting its sequestration at high-affinity sites or creating a stoichiometric imbalance that disturbs maintenance complex assembly. Perturbation of other components of the DNA methylation machinery may have also contributed. We therefore analyzed gene expression of other key regulators of DNA methylation^1^, including *Uhrf1*, *Zfp57*, *Zfp445*, *Dnmt3a*, *Dnmt3b*, *Dnmt3l*, *Tet1*, *Tet2* and *Tet3*. Notably, *Dnmt3b* and *Dnmt3l* were reduced in *Dnmt1^tet/tet^* mESCs to 0.4- and 0.3-fold of wild-type levels, respectively, whereas the others were unchanged (Figure 1B; Data S1). Correspondingly, DNMT3B protein was diminished in *Dnmt1^tet/tet^* mESCs, down to 0.1-fold relative to wild-type mESCs (Figures 1C–D, S1; Data S2). DNMT3L protein was not assessed due to poor antibody specificity, but its reduced gene expression in *Dnmt1^tet/tet^* mESCs prompted us to also examine DNMT3A protein, as Veland et al. demonstrated that DNMT3L stabilizes DNMT3A and protects it from degradation in mESCs^13^. Supporting this rationale, DNMT3A protein was reduced in *Dnmt1^tet/tet^* mESCs, to 0.01-fold for DNMT3A1 (long isoform) and 0.4-fold for DNMT3A2 (short isoform) (Figures 1C–D, S1; Data S2), despite unchanged gene expression. The DNMT3A1 fold change did not reach statistical significance, presumably due to variable western blot band intensities in wild-type samples. We suspected the strongest wild-type band to be an outlier, but Dixon’s Q test did not identify it as such (p-value = 0.1009). Nevertheless, *Dnmt1^tet/tet^* samples displayed a clear absence of DNMT3A1 bands. DNMT3A and DNMT3B mediate de novo methylation^1^, but they have also been shown to contribute to the maintenance of pre-existing methylation patterns^5–7^. Their reduction, together with diminished support from DNMT3L, may therefore compromise both de novo methylation and maintenance fidelity in *Dnmt1^tet/tet^* mESCs, thereby contributing to global hypomethylation. While stable de novo methylation would not typically be expected in mESC cultures under normal conditions, transient de novo activity during cyclic demethylation-remethylation events is essential^39^.

To our knowledge, DNMT1 overexpression leading to reduced DNMT3A, DNMT3B, or DNMT3L has not been previously reported. In cancer—the most studied context of DNMT1 overexpression—DNMT3A and DNMT3B are typically elevated as well^24^. However, DNMT overexpression in cancer is likely a downstream consequence of other oncogenic programs, whereas in *Dnmt1^tet/tet^* mESCs, DNMT1 is directly overexpressed in otherwise healthy cells, making it the primary driver of the observed effects. Non-transformed cells may actively compensate for DNMT1 excess in ways that cancer cells, already burdened by widespread regulatory disruption, cannot. Isolating the consequences of direct DNMT1 overexpression in mESCs therefore provides a useful framework for disentangling the contributions of DNMT1 itself from the secondary effects of oncogenic signaling in cancer.

Taken together, these results call attention to the cooperativity among DNMT family members, whereby DNA methylation regulation depends not only on the activity of individual DNMTs but also on their balanced dosage and coordinated interplay, with perturbations in one member reverberating across the family. We observed global hypomethylation alongside reduced DNMT3A and DNMT3B (and plausibly DNMT3L, confirmed only at the transcript level) in *Dnmt1^tet/tet^* mESCs, suggesting DNMT1 overexpression may paradoxically compromise its maintenance efficiency rather than reinforcing it, while also perturbing the broader DNA methylation machinery. Paradoxical or compensatory outcomes of this kind are not unprecedented; Kafetzopoulos et al. showed that complete DNMT1 loss in colorectal cancer cells does not produce uniform hypomethylation but instead generates a mixed landscape: most late-replicating, heterochromatic PMDs become hypomethylated, whereas a subset of H3K9me3-marked domains gain methylation through DNMT3A recruitment following chromatin reconfiguration^40^. This highlights how disrupting one DNMT can provoke domain-specific compensation by others, reinforcing that methylation homeostasis reflects coordinated, context-dependent interplay rather than isolated enzyme function. However, global hypomethylation in *Dnmt1^tet/tet^* mESCs was relatively modest (8% loss), raising the possibility that excess DNMT1 may have also stimulated its non-canonical de novo activity^2–4^, producing a composite effect in which hypomethylation was offset by localized hypermethylation. Resolving this duality will require more granular, locus-specific methylation analyses.

### Genome-wide hypomethylation, local hypermethylation and higher CpG-to-CpG variability in *Dnmt1^tet/tet^* mESCs

To dissect the impact of DNMT1 overexpression on DNA methylation with higher resolution, we identified differentially methylated regions (DMRs) in *Dnmt1^tet/tet^* versus wild-type mESCs. DMRs were called using the DSS R package^41^, which generates regions of differential methylation with variable lengths and CpG densities, while accounting for biological and sequencing coverage variability across samples. We defined DMRs as regions at least 100 bp in length and containing at least five CpG sites, in which at least 50% of CpGs exhibited a significant (adjusted p-value ≤ 0.01) methylation change of at least 10%, and the regional mean methylation change was also at least 10%. Hypomethylated DMRs were vastly more numerous than hypermethylated DMRs (81,144 versus 4,303) (Figure 2A) and tended to be longer (mean length of 1,238 bp versus 470 bp, median length of 794 bp versus 436 bp) (Figure 2B), aligning with DNA methylation being globally reduced (Figure 1E). Moreover, hypomethylated DMRs exhibited lower CpG densities and were located farther from transcription start sites (TSSs) compared to hypermethylated DMRs (Figure 2C–D). Reflecting these properties, hypermethylated DMRs were concentrated in promoter regions nearest to TSSs (core and proximal promoters) and in CpG islands (CpG-dense regions), with nearly equal representation of hypermethylated and hypomethylated DMRs in core promoters and striking overrepresentation of hypermethylated DMRs in CpG islands (Figure 2E; Table S1). This localized prominence of hypermethylated DMRs in *Dnmt1^tet/tet^* mESCs is consistent with previous studies linking DNMT1 overexpression to promoter and CpG island hypermethylation^24,42–44^.

**Figure 2.**
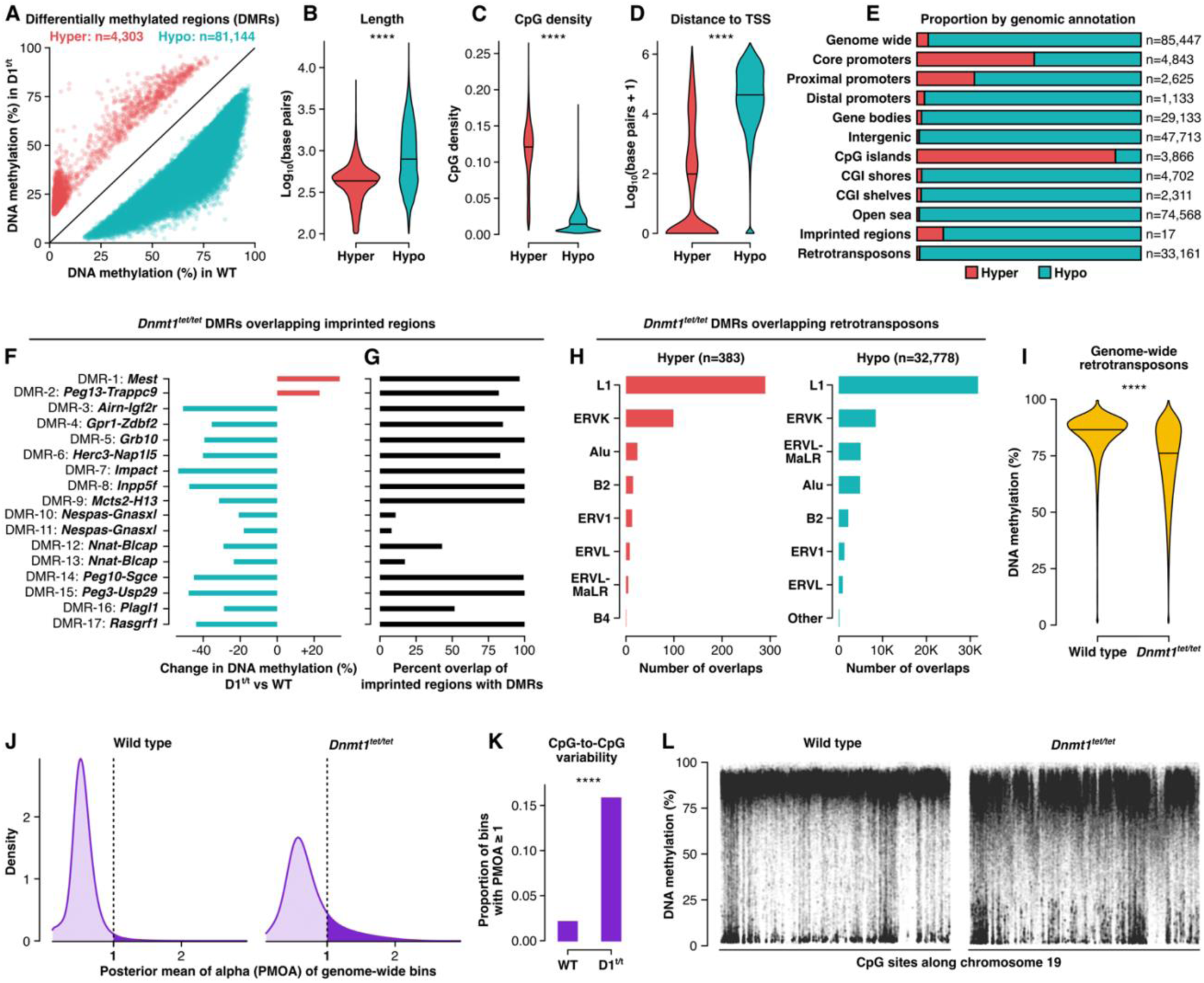
Genome-wide hypomethylation, local hypermethylation and higher CpG-to-CpG variability in *Dnmt1^tet/tet^* mESCs. **(A)** Differentially methylated regions (DMRs) identified in *Dnmt1^tet/tet^* versus wild-type mESCs using EM-seq (n=3 per condition) and the DSS R package. All DMRs are ≥ 100 bp in length, contain ≥ 5 CpGs, have ≥ 50% of CpGs exhibiting a significant (adjusted p-value ≤ 0.01) methylation change ≥ 10%, and have a regional mean methylation change ≥ 10%. Hyper : hypermethylated, hypo : hypomethylated. **(B)** Length in base pairs of *Dnmt1^tet/tet^* hyper and hypo DMRs. **(C)** CpG density of *Dnmt1^tet/tet^* hyper and hypo DMRs measured using all genomic CpGs from the mm10 reference genome (not just sequenced CpGs). **(D)** Distance in base pairs between nearest transcription start site and *Dnmt1^tet/tet^* hyper and hypo DMRs. Statistical comparisons in **B**, **C**, and **D** were conducted using the Kruskal-Wallis test. **** p-value ≤ 0.0001. **(E)** Proportion of *Dnmt1^tet/tet^* hyper and hypo DMRs by genomic annotation. *See also Table S1*. **(F)** Change in DNA methylation in *Dnmt1^tet/tet^* versus wild-type mESCs for DMRs that overlap imprinted regions. **(G)** Percent overlap of imprinted regions with *Dnmt1*^tet/tet^ DMRs. **(H)** Number of *Dnmt1^tet/tet^* hyper and hypo DMRs that overlap loci of specific retrotransposon families. **(I)** DNA methylation levels of genome-wide retrotransposons measured by EM-seq in wild-type and *Dnmt1^tet/tet^* mESCs (n=3 per condition). Statistical comparison was conducted using the Kruskal-Wallis test. **(J)** Distribution of posterior mean of alpha (PMOA) values of genome-wide bins of 101 CpGs in wild-type and *Dnmt1^tet/tet^* mESCs calculated using the MethylSeekR R package. Unimodal distribution with a peak < 1 indicates a polarized methylome with low CpG-to-CpG variability and bimodal distribution with a second peak ≥ 1 indicates the presence of partially methylated domains. PMOA value ≥ 1 indicates CpG-to-CpG variability. **(K)** Proportion of genomic bins with a PMOA value ≥ 1 in wild-type and *Dnmt1^tet/tet^* mESCs. Statistical comparison was conducted using Pearson’s chi-squared test. **(L)** DNA methylation levels of single-CpG sites along chromosome 19 in wild-type and *Dnmt1^tet/tet^* mESCs, demonstrating the higher CpG-to-CpG variability in the *Dnmt1^tet/tet^* methylome.

We next examined DNA methylation of imprinted regions and retrotransposons more closely, given DNMT1’s established role in genomic imprinting maintenance^45^ and Haggerty et al. reporting its de novo activity at retrotransposons in mESCs^3^. Imprinted regions overlapped with many more hypomethylated than hypermethylated DMRs (15 versus 2), with most affected regions nearly fully encompassed by a DMR (Figure 2E–G; Table S1). Hypomethylation of entire imprinted regions in *Dnmt1^tet/tet^* mESCs suggests that the protein complex responsible for maintaining genomic imprints, which includes DNMT1^46^, could have been defective. Seeing as expression levels of *Uhrf1*, *Zfp57* and *Zfp445*—other components of this complex^46^—were unchanged in *Dnmt1^tet/tet^* mESCs (Figure 1B; Data S1), it is unlikely that a defect in imprinting maintenance would arise from a shortage of components; rather, the stoichiometric balance needed for proper complex assembly may have been disrupted by excess DNMT1. Retrotransposons likewise overlapped predominantly with hypomethylated DMRs than with hypermethylated DMRs (32,778 versus 383), most frequently at LINE-1 (long interspersed nuclear element-1) and ERVK (endogenous retrovirus group K) loci (Figure 2E & H; Table S1). Their mean methylation loss of 12% (Figure 2I) exceeded the mean genome-wide loss of 8% (Figure 1E); mean retrotransposon methylation was 83% in wild-type and 71% in *Dnmt1^tet/tet^* mESCs (Figure 2I). The scarcity of hypermethylation at retrotransposons in *Dnmt1^tet/tet^* mESCs likely reflects their already high baseline methylation levels in wild-type cells, limiting further gains. In contrast, in the study by Haggerty et al., global hypomethylation was first induced in mESCs by depleting all DNMTs before only re-expressing DNMT1 to assess its de novo activity^3^, thereby creating potential for de novo methylation at retrotransposons.

Retrotransposon hypomethylation in *Dnmt1^tet/tet^* mESCs could result from excess DNMT1 causing it to preferentially localize to other genomic sites, such as promoters and CpG islands, sequestering it away from retrotransposons. Moreover, reduced DNMT3A and DNMT3B protein levels in *Dnmt1^tet/tet^* mESCs (Figures 1C–D, S1; Data S2) may have contributed to hypomethylation of both retrotransposons and imprinted regions, as they have been shown to support methylation maintenance at these loci^5–7^.

Given that *Dnmt1^tet/tet^* mESCs exhibited global loss of DNA methylation (Figures 1E, 2A), we questioned whether partially methylated domains (PMDs) could have emerged. PMDs are large genomic regions characterized by reduced average methylation relative to background levels and high methylation variability between adjacent CpG sites (i.e. CpG-to-CpG variability)^47^. PMDs are observed in some somatic cell types^48^ and in cancer^49,50^ but are normally absent in ESCs^48^. To detect the presence of PMDs, we used the MethylSeekR R package that calculates the posterior mean of alpha (PMOA) across genomic bins of 101 CpGs^48^. A unimodal PMOA distribution with a peak below 1 is indicative of a methylome without PMDs, whereas a bimodal distribution with a secondary peak above 1 is indicative of a methylome with PMDs. In methylomes that lack PMDs but exhibit high variability, a notable fraction of PMOA values (10-20%) may exceed 1, with the overall distribution remaining unimodal. Both wild-type and *Dnmt1^tet/tet^* methylomes exhibited unimodal PMOA distributions with peaks below 1, meaning the presence of PMDs was not indicated (Figure 2J). However, the *Dnmt1^tet/tet^* methylome had a greater proportion of bins with PMOA values above 1 (16% versus 2%) (Figure 2K), reflecting higher CpG-to-CpG variability. Single-CpG methylation levels were plotted along chromosome 19, visually demonstrating the more variable methylome of *Dnmt1^tet/tet^* mESCs in comparison to the more polarized methylome of wild-type mESCs (Figure 2L). This increase in CpG-to-CpG variability, together with global hypomethylation, further supports compromised methylation maintenance in *Dnmt1^tet/tet^* mESCs. During maintenance, a process known as neighbour-guided correction, proposed by Wang et al., is thought to leverage the methylation status of adjacent CpGs to ensure fidelity^51^. Loss of methylation due to faulty deposition could undermine neighbour-guided correction, increasing CpG-to-CpG variability. Increased variability, in turn, may further impair correction fidelity, exacerbating methylation loss. Such a cascade could have been triggered in *Dnmt1^tet/tet^* mESCs by the aberrant dosage of DNMTs (Figures 1C–D, S1; Data S2).

To summarize, DNMT1 overexpression in mESCs led to genome-wide DNA hypomethylation—notably affecting many transposable elements and imprinted regions—alongside localized hypermethylation at promoters and CpG islands, most likely arising from reduced maintenance efficiency together with increased de novo activity. Moreover, the *Dnmt1^tet/tet^* methylome exhibited higher CpG-to-CpG variability, suggesting impaired neighbour-guided correction during maintenance, which could have accelerated methylation loss. These findings have compelling parallels with cancer, which has been associated with DNMT1 overexpression^24,42–44^. Similarly to *Dnmt1^tet/tet^* mESCs, cancer methylomes often exhibit a combination of genome-wide hypomethylation, promoter CpG island hypermethylation, de-repression of transposable elements, loss of genomic imprints and high CpG-to-CpG variability^25,26^. While PMDs did not emerge in the *Dnmt1^tet/tet^* methylome, its increased CpG-to-CpG variability could point to a consequence of defective methylation maintenance that, in cancer, culminates in the disproportionate methylation loss in PMDs versus non-PMD regions^52^. Because PMDs are intrinsically highly variable^47^ and would therefore already be maintained with reduced fidelity, their response to defective methylation maintenance could be magnified, further increasing their variability and accelerating their erosion. In contrast, non-PMD regions, or methylomes with low baseline variability such as in ESCs^48^, may absorb the impact more evenly and effectively, resulting in slower erosion and avoiding domain-scale methylation collapse. Ultimately, these findings reveal that DNMT1 overexpression alone is sufficient to mimic several cancer-associated DNA methylation features in mESCs.

### Promoter DMRs in *Dnmt1^tet/tet^* mESCs align with the pathological potential of DNMT1 overexpression

DNMT1 overexpression has been studied abundantly in the context of cancer^24,42–44^ but has also been linked to other diseases, including cardiomyopathy^53^, schizophrenia^54^, epilepsy^55^ and bipolar disorder^56^. DNMT1 overexpression also has severe developmental consequences; Biniszkiewicz et al. reported midgestational lethality in mouse embryos^37^. Embryonic survival until midgestation may reflect the greater plasticity of earlier developmental stages^57^, in contrast to midgestation, a critical phase of organogenesis and tissue differentiation^58^ demanding stricter regulatory control. Consistent with this idea, D’Aiuto et al. demonstrated that *Dnmt1^tet/tet^* mESCs form morphologically normal embryoid bodies, indicative of relatively stable early differentiation, but later generate neurons with abnormal dendritic arborization and branching^34^. Compellingly, we found that hypermethylated promoter DMRs in *Dnmt1^tet/tet^* mESCs were associated with genes involved in axon guidance, developmental signaling (Hippo, Wnt, pluripotency) and cancer (breast, gastric, hepatocellular carcinoma), and hypomethylated promoter DMRs with genes involved in multiple neuronal pathways (neuroactive ligand signaling and receptor interaction, glutamatergic synapse, axon guidance) and cardiomyopathy (arrhythmogenic right ventricular and dilated cardiomyopathy) (Figure 3A; Data S3, S4). These findings suggest that *Dnmt1^tet/tet^* mESCs harbor promoter methylation alterations that may foreshadow deviations from normal developmental trajectories and provide mechanistic insight into many relevant diseases.

**Figure 3.**
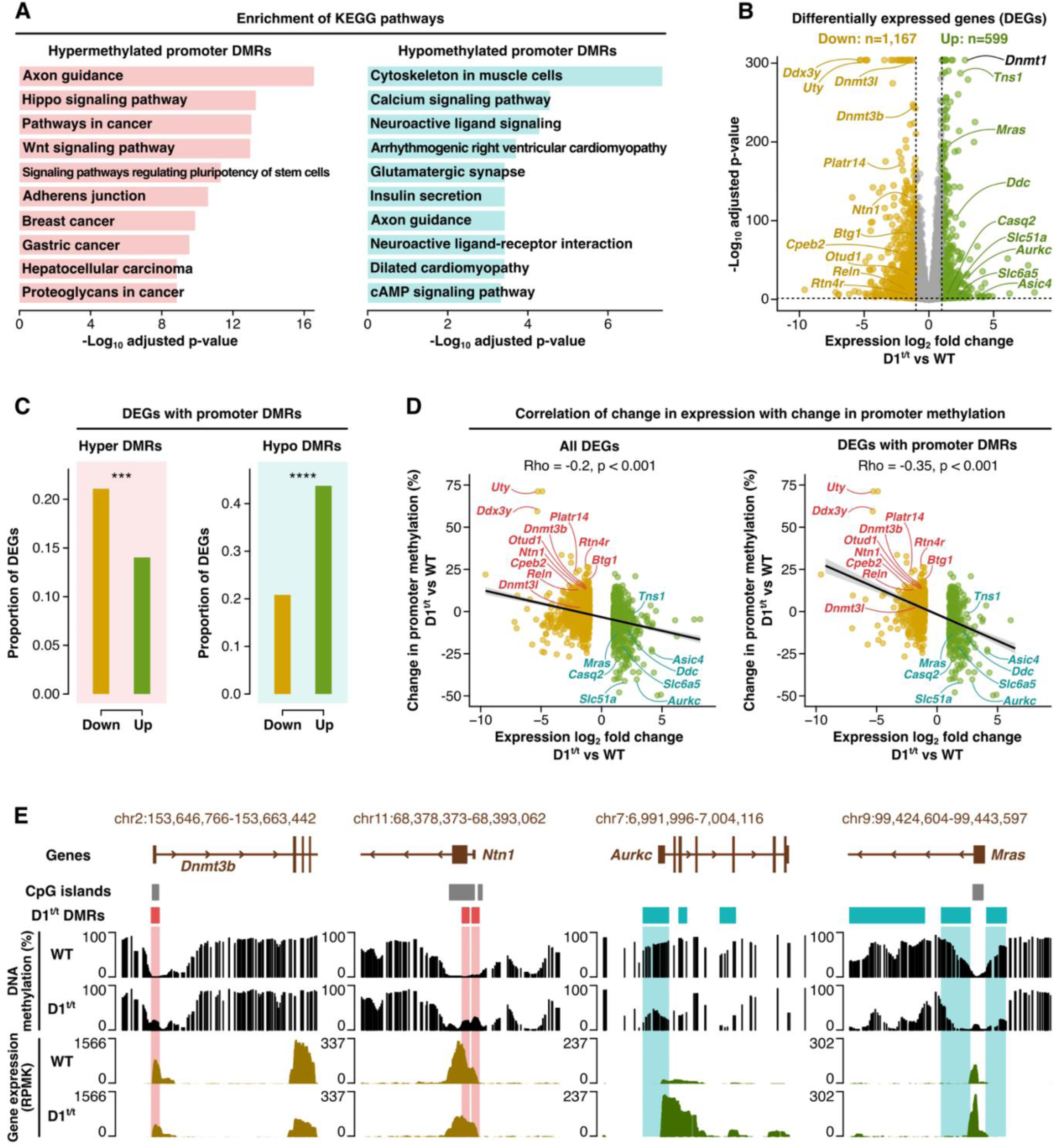
Promoter DMRs in *Dnmt1^tet/tet^* mESCs align with the pathological potential of DNMT1 overexpression. **(A)** Enrichment of KEGG pathways in genes linked to hypermethylated (hyper) and hypomethylated (hypo) *Dnmt1^tet/tet^* DMRs in promoters. Enrichment was performed with the clusterProfiler R package against a background universe of all genes with available promoter methylation data. *See also Data S3 and S4*. **(B)** Differentially expressed genes (DEGs) in *Dnmt1^tet/tet^* versus wild-type mESCs measured by mRNA-seq (n=3 per condition) and quantified using the DESeq2 R package. Only genes with available promoter DNA methylation levels were included. Statistical significance was defined by a log_2_ fold change ≤ -1 or ≥ 1 and an adjusted p-value ≤ 0.05. *See also Data S1*. **(C)** Proportion of *Dnmt1^tet/tet^* downregulated (down) and upregulated (up) DEGs containing hyper or hypo DMRs in their promoters. Statistical comparison was conducted using Pearson’s chi-squared test. **** p-value ≤ 0.0001, *** p-value ≤ 0.001. **(D)** Spearman’s correlation of gene expression log_2_ fold change with promoter DNA methylation change in *Dnmt1*^tet/tet^ versus wild-type mESCs for all DEGs and only for DEGs with promoter DMRs. **(E)** Genomic tracks showing examples of DEGs with promoter DMRs.

We next investigated how promoter methylation alterations related to changes in gene expression. Quantifying expression for all genes with available promoter methylation data, we identified 1,167 downregulated and 599 upregulated differentially expressed genes (DEGs) in *Dnmt1^tet/tet^* relative to wild-type mESCs (Figure 3B; Data S1). Consistent with the generally repressive role of promoter methylation^1^, hypermethylated promoter DMRs were more prevalent among downregulated than upregulated DEGs (Figure 3C). Conversely, hypomethylated promoter DMRs were more prevalent among upregulated than downregulated DEGs (Figure 3C), reflecting the activating potential of reduced promoter methylation^1^. Furthermore, change in promoter methylation levels negatively correlated with differential gene expression (rho = -0.2), with a stronger correlation (rho = -0.35) when limiting the analysis to DEGs with promoter DMRs (Figure 3D–E). Among the upregulated DEGs with hypomethylated promoter DMRs were *Slc6a5*^59^, *Asic4*^60^ and *Ddc*^61^, involved in neurotransmission and synaptic regulation, and *Tns1*^62^, *Casq2*^63^ and *Mras*^64^, which contribute to cardiac development and function (Figure 3D). Downregulated DEGs with hypermethylated promoter DMRs included *Cpeb2*^65^, *Otud1*^66^ and *Btg1*^67^, which have been linked to cancer, and *Rtn4r*^68^, *Reln*^69^ and *Ntn1*^70^, critical for neural development and function (Figure 3D). Notably, *Rtn4r* is located within the 22q11.2 chromosomal region, a well-established hotspot associated with schizophrenia and autism^68^. *Dnmt3b* and *Dnmt3l* were also included among the downregulated DEGs with hypermethylated promoter DMRs (Figure 3D). DNMT3B showing promoter hypermethylation together with lower expression and protein levels (Figures 1C–D, 3D, S1; Data S2) follows the classic epigenetic dogma, suggesting it was directly targeted for repression in response to DNMT1 overexpression rather than through an indirect compensatory mechanism. Although DNMT3L protein could not be verified by western blot due to antibodies lacking specificity, hypermethylation in its promoter and its downregulated expression (Figure 3D) point to a similar effect, strengthening our hypothesis that reduced DNMT3L protein in *Dnmt1^tet/tet^* mESCs is to blame for DNMT3A having lower protein levels despite unchanged expression (Figures 1B–D, S1; Data S1, S2), given DNMT3L’s role in stabilizing DNMT3A in mESCs^13^. Overall, the antagonistic relationship between promoter methylation and gene expression in *Dnmt1^tet/tet^* mESCs aligns with core principles of epigenetic regulation^1^. However, the modest strength of their correlation underscores how promoter methylation does not dictate transcriptional output in isolation but instead operates within broader regulatory contexts involving biological and environmental cues, other epigenetic marks, large-scale chromatin architecture and transcription factor availability. Differential gene expression in *Dnmt1^tet/tet^* mESCs may also stem from DNA methylation-independent mechanisms such as cascading effects, mitigation by cellular plasticity and non-catalytic DNMT1 activity, which primarily involves protein-protein interactions with its N-terminal domains^71^.

In brief, *Dnmt1^tet/tet^* promoter DMRs were associated with cancer, cardiac disease, neuronal and developmental pathways, consistent with the broad pathological potential of DNMT1 overexpression. Moreover, promoter methylation and gene expression changes were correlated, but the modest strength of their relationship points to additional mechanistic layers. Dissecting these layers will be essential to clarify how DNMT1 overexpression reshapes regulatory networks, with *Dnmt1^tet/tet^* mESCs proving to be a developmentally and disease-relevant model.

### Binding sites of key transcriptional regulators intersect with *Dnmt1^tet/tet^* promoter DMRs

To explore how the regulatory impact of promoter methylation alterations in *Dnmt1^tet/tet^* mESCs may involve additional mechanistic layers, we intersected promoter DMRs with binding sites of transcriptional regulators obtained from the mouse ReMap dataset catalog^72^ and performed enrichment analysis.

*Dnmt1^tet/tet^* hypermethylated promoter DMRs notably overlapped with binding sites of HDAC1 (histone deacetylase 1) and E2F1 (E2F transcription factor 1) (Figure 4A; Data S5). Fuks et al. previously reported that HDAC1 has the ability to bind DNMT1^15^, and prior work by Robertson et al. showed that DNMT1 forms a repressive complex with HDAC1, E2F1 and Rb (retinoblastoma protein) that targets E2F-responsive genes in HeLa cells derived from human cervical cancer^73^. HDAC1 promotes chromatin compaction by deacetylating histone tails, thereby participating in transcriptional repression^74^, while E2F1 is a transcriptional activator that regulates cell cycle progression, DNA replication and repair, apoptosis and differentiation^75^. E2F1 functions are context-dependent, promoting either cell survival or cell death, interacting with Rb to restrain inappropriate cell cycle entry^76^. Although we did not observe significant overlap between *Dnmt1^tet/tet^* hypermethylated promoter DMRs and Rb binding sites (Data S5), it cannot be excluded given dataset limitations that could have reduced statistical power: the mouse ReMap catalog contains three Rb datasets with a median of only 1,233 binding sites, compared with five datasets for E2F1 with a median of 7,824 sites and 19 datasets for HDAC1 with a median of 17,398 sites. Increased availability of DNMT1 protein in *Dnmt1^tet/tet^* mESCs could raise the likelihood of the DNMT1-HDAC1-E2F1-Rb complex assembling, leading to locus-specific de novo methylation at E2F1 binding sites and abnormal gene repression. Having observed that *Dnmt1^tet/tet^* hypermethylated DMRs were highly concentrated in CpG islands (Figure 2D; Table S1) further supports this possibility given that E2F1 binding sites are commonly located within or near CpG islands^77^. Regional co-occurrence analysis of CpG islands, E2F1 binding sites and *Dnmt1^tet/tet^* hypermethylated promoter DMRs revealed that hypermethylation indeed occurred precisely within CpG islands overlapping E2F1 binding sites (Figure 4B). For example, this was observed at the promoters of *Ccne1*, a key regulator of G1/S phase transition in the cell cycle^78^, and *Casp9*, the primary initiator of the intrinsic apoptotic pathway^79^ (Figure 4C). Also noteworthy were *Dnmt1^tet/tet^* hypermethylated promoter DMRs being enriched at binding sites of CREBBP (CREB-binding protein) and TEAD1 (TEA Domain Transcription Factor 1) (Figure 4A; Data S5). CREBBP promotes chromatin accessibility and transcriptional activation through histone acetylation^80^; hypermethylation of its binding sites may counteract this activity, as DNA methylation generally antagonizes histone acetylation in promoters^81^. TEAD1, a transcriptional activator in the Hippo signaling pathway, binds DNA with YAP (yes-associated protein), TAZ (WW domain-containing transcription regulator 1) and additional partners^82^. Multi-protein complexes of this kind require ample chromatin accessibility and flexibility for effective binding, which could be sterically hindered by a local increase in DNA methylation. Collectively, binding sites of these transcriptional regulators overlapping with *Dnmt1^tet/tet^* hypermethylated promoter DMRs highlight potential canonical relationships where promoter hypermethylation could lead to transcriptional repression.

**Figure 4.**
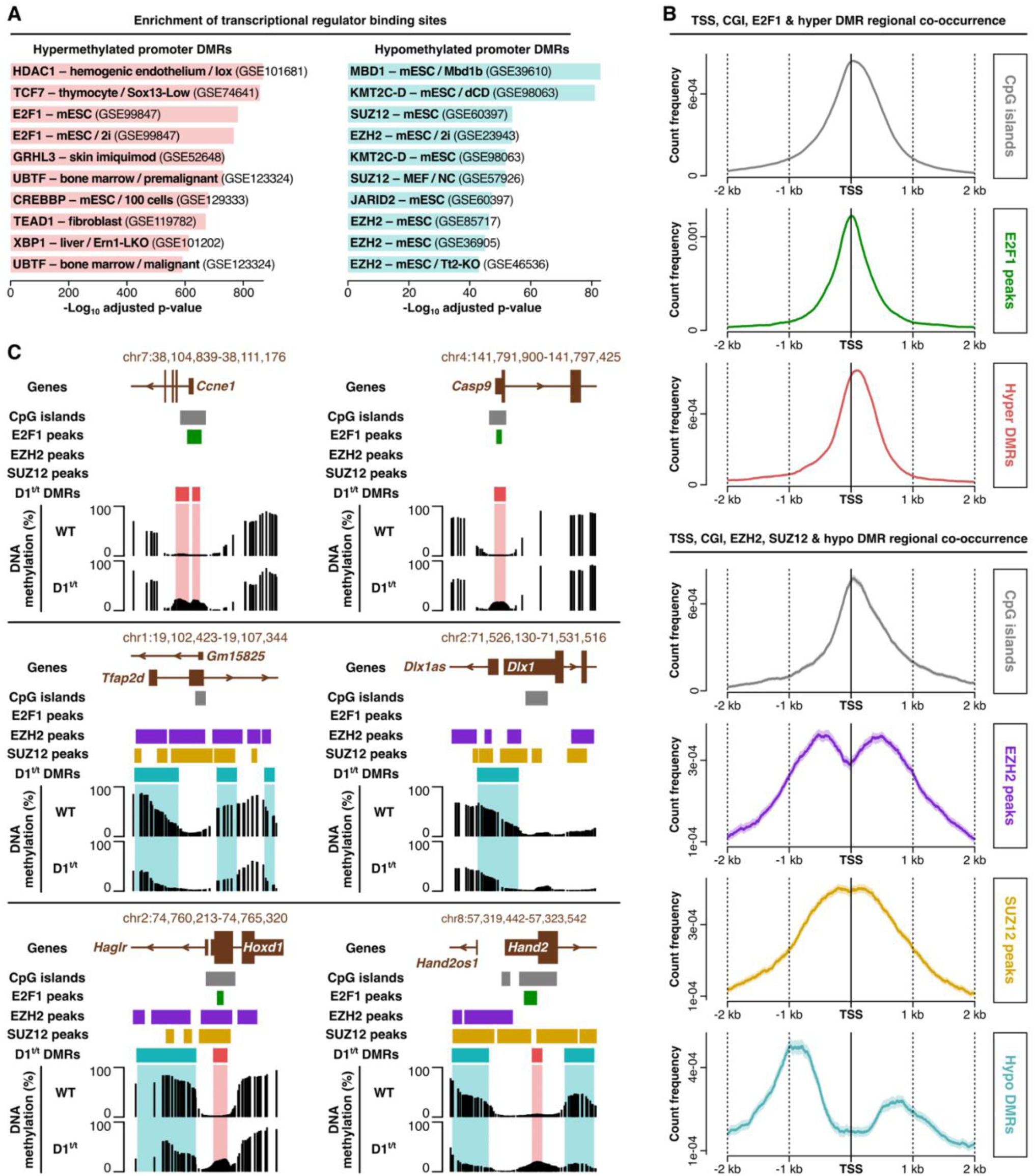
Binding sites of key transcriptional regulators intersect with *Dnmt1^tet/tet^* promoter DMRs. **(A)** Enrichment of transcriptional regulator binding sites that intersect hypermethylated (hyper) and hypomethylated (hypo) *Dnmt1^tet/tet^* DMRs in promoters. Binding sites were obtained from the mouse ReMap dataset catalog. Enrichment was performed with the ReMapEnrich R package using 5000 permutation shuffles against a background universe of all promoter regions with available methylation data. *See also Data S5 and S6*. **(B)** Regional co-occurrence of transcription start sites (TSS), CpG islands (CGI), transcriptional regulator binding sites, and *Dnmt1^tet/tet^* hypo or hyper DMRs. Only datasets of transcriptional regulator binding sites identified as significantly enriched in **A** and generated from wild-type mESCs cultured in serum were used. **(C)** Genomic tracks showing examples of regions that were examined in **B**.

On the other hand, overlaps between *Dnmt1^tet/tet^* hypomethylated promoter DMRs and transcriptional regulator binding sites provide insight into why promoter methylation and gene expression alterations may not always align in a predictable manner. Specifically, enrichment was observed at binding sites of MBD1 (methyl-CpG binding domain protein 1), KMT2C–D (histone lysine methyltransferases 2C and 2D) and PRC2 (polycomb repressive complex 2) components SUZ12, EZH2 and JARID2 (Figure 4A; Data S6). While hypomethylation at MBD1 binding sites could activate transcription since MBD1 normally binds methylated CpGs to stabilize repressive chromatin^83^, hypomethylation at KMT2C–D and PRC2 binding sites could drive activating, repressive and neutral effects. In mESCs, many developmental promoters are bivalent, marked simultaneously by activating and repressive histone modifications, which creates a poised chromatin state that is thought to enable rapid activation or repression throughout development^84^. Classically, bivalent promoters carry activating H3K4me3 (histone H3 lysine 4 trimethylation) together with repressive H3K27me3 (histone H3 lysine 27 trimethylation), with PRC2 catalyzing the latter^84^. More recently, trivalent states including H3K4me1 (histone H3 lysine 4 monomethylation)^85^, catalyzed by KMT2C–D^86^, have also been described. These promoters typically harbor unmethylated CpG islands flanked by methylated shores; methylation of CpG island shores is believed to serve as an epigenetic boundary limiting the spread of histone modifications^87,88^. Their hypomethylation could thus allow spreading of activating, repressive or both types of histone modifications, thereby resulting in transcriptional activation, repression or neutral outcomes. Supporting these possibilities, regional co-occurrence of CpG islands, binding sites of core PRC2 components EZH2 and SUZ12, and *Dnmt1^tet/tet^* hypomethylated promoter DMRs showed that hypomethylation predominantly flanked the CpG islands (Figure 4B). For example, this was observed at the promoters of *Tfap2d*^89^ and *Dlx1*^90^, two developmental transcription factors (Figure 4C). This resonates with having detected DNMT3A1 protein in wild-type but not in *Dnmt1^tet/tet^* mESCs (Figure 1C), as Manzo et al. identified DNMT3A1 as a principal mediator of CpG island shore methylation^87^. Interestingly, in some regions we observed co-occurrence of CpG islands, binding sites of EZH2, SUZ12 and E2F1, together with *Dnmt1^tet/tet^* hypomethylated and hypermethylated promoter DMRs, reinforcing that promoter methylation alterations may yield complex transcriptional outcomes. For example, promoters of *Hoxd1*^91^ and *Hand2*^92^, also developmental transcription factors, exhibited such characteristics (Figure 4C).

In summary, by overlapping *Dnmt1^tet/tet^* promoter DMRs with binding sites of transcriptional regulators from public datasets, we uncovered potential mechanisms implicated in their regulatory impact. These included canonical relationships, such as promoter hypermethylation coinciding with E2F1, TEAD1, HDAC1 and CREBBP, and hypomethylation with MBD1. More complex relationships were also implied, such as hypomethylation of PRC2 binding sites as well as concomitant hypermethylation and hypomethylation in target promoters of both E2F1 and PRC2. However, given the context-dependent functions of these transcriptional regulators, ChIP-seq (chromatin immunoprecipitation sequencing) validation in more appropriate developmental and disease settings will be needed to ascertain their specific roles in rewiring regulatory networks upon DNMT1 overexpression.

### Hypermethylation in *Dnmt1^tet/tet^* mESCs partially persists following DNMT1 depletion

To counteract cancer-associated hypermethylation, pan-DNMT inhibitors (e.g. azacitidine, decitabine) are approved for clinical use^29^ and selective DNMT1 inhibitors (e.g. GSK3685032, GSK3482364) are being studied in pre-clinical models^30^. This prompted us to investigate whether hypermethylation observed in *Dnmt1^tet/tet^* mESCs could be effectively erased following DNMT1 depletion via doxycycline (dox) treatment (Figure 1A); after three days of dox treatment, DNMT1 protein becomes undetectable^31^, and after six days, almost all DNA methylation is lost through passive demethylation^33^. Consistent with this, six days of dox treatment silenced *Dnmt1* expression (Figure 1B; Data S1), rendered DNMT1 protein undetectable (Figures 1C–D, S1; Data S2) and reduced global DNA methylation levels to 4% (Figure 1E).

Despite drastic global hypomethylation in dox-treated *Dnmt1^tet/tet^* mESCs, methylation levels of *Dnmt1^tet/tet^* hypermethylated DMRs (Figure 2A) remained elevated overall (Figure 5A), with 1,708 of them showing a persistent increase of at least 10% (Figure 5B; Table S1). While the majority (n = 1,146) did exhibit some methylation loss in dox-treated *Dnmt1^tet/tet^* mESCs, many (n = 475) maintained similar levels, and some (n = 87) even showed further gains (Figure 5C). *Dnmt1^tet/tet^* hypermethylated DMRs in core promoters and CpG islands were most strongly associated with persistent hypermethylation in dox-treated *Dnmt1^tet/tet^* mESCs (Figure 5D; Table S1), notably affecting cancer-related genes (*Cpeb2*^65^, *Otud1*^66^, *Tbx15*^93^, *Ppp2r5e*^94^) as well as nervous system-related genes (*Rtn4r*^68^, *Reln*^69^, *Kcnma1*^95^, *Rest*^96^) (Figure 5E). Persistent hypermethylation in dox-treated *Dnmt1^tet/tet^* mESCs strongly suggests that DNMT3A and DNMT3B maintained hypermethylation in the absence of DNMT1, contributed to its initial deposition or both, thereby rendering DNMT1 depletion alone insufficient for its complete erasure. Although DNMT3A and DNMT3B levels were reduced in *Dnmt1^tet/tet^* and dox-treated *Dnmt1^tet/tet^* mESCs, their presence remained detectable in both conditions (Figures 1C–D, S1; Data S2). Clarifying their roles will require further investigation, for example via knockdown experiments using small interfering RNA—specific inhibitors for DNMT3A and DNMT3B are not yet available.

**Figure 5.**
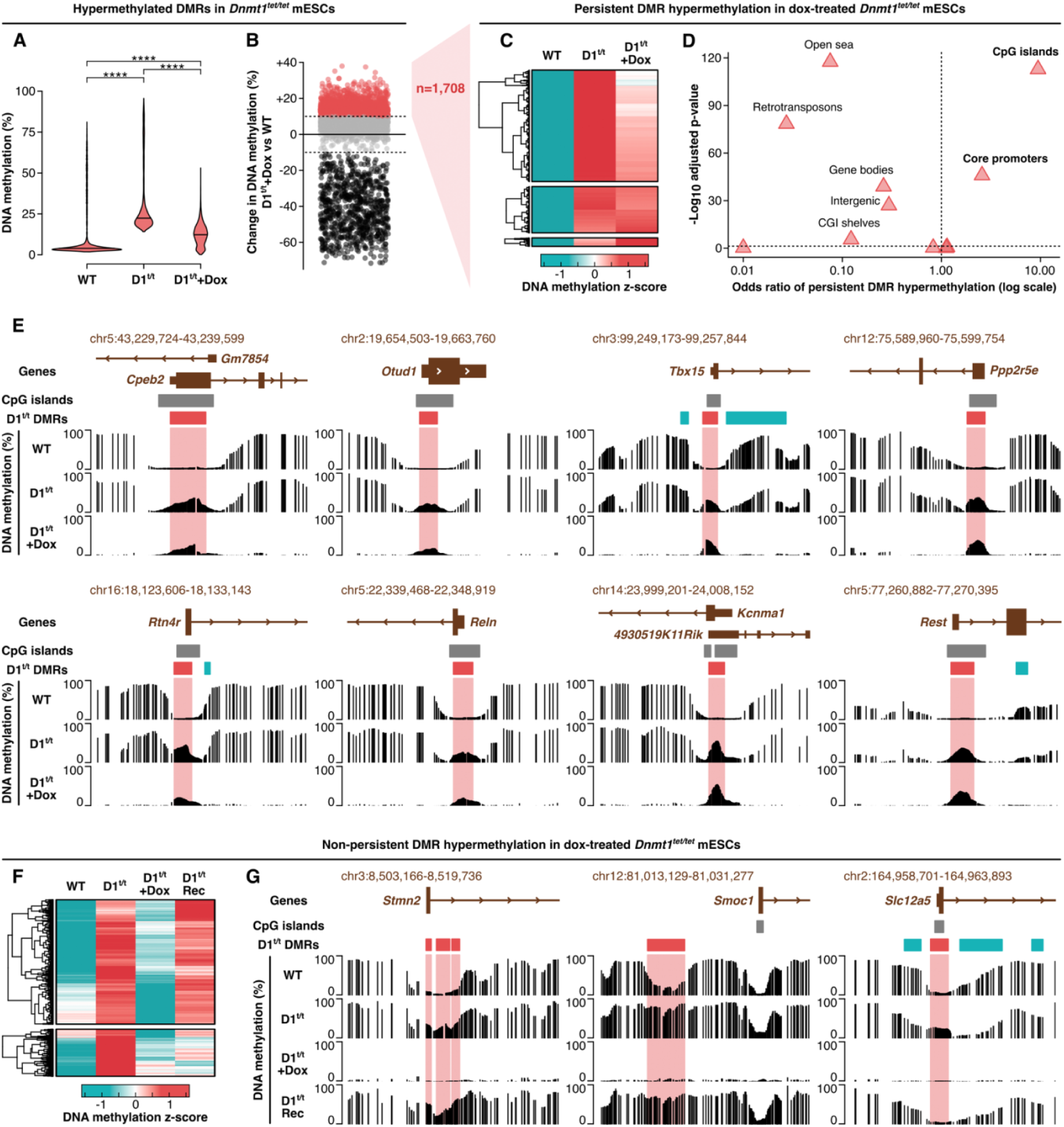
Hypermethylation in *Dnmt1^tet/tet^* mESCs partially persists following DNMT1 depletion. **(A)** DNA methylation levels of *Dnmt1^tet/tet^* hypermethylated DMRs in wild-type, *Dnmt1^tet/tet^* and doxycycline-treated *Dnmt1^tet/tet^* mESCs measured by EM-seq (n=3 per condition). Statistical comparisons were conducted using the pairwise Wilcoxon test with Benjamini-Hochberg p-value adjustment. **** p-value ≤ 0.0001. **(B)** Change in DNA methylation in doxycycline-treated *Dnmt1^tet/tet^* versus wild-type mESCs for *Dnmt1^tet/tet^* hypermethylated DMRs. **(C)** Z-score normalization of DNA methylation levels in wild-type, *Dnmt1^tet/tet^* and doxycycline-treated *Dnmt1^tet/tet^* mESCs for *Dnmt1^tet/tet^* hypermethylated DMRs that showed ≥ 10% increase in DNA methylation in doxycycline-treated *Dnmt1^tet/tet^* versus wild-type mESCs in **B** i.e. persistent DMR hypermethylation. **(D)** Odds ratios showing the strength of association between persistent DMR hypermethylation and specific genomic annotations. An odds ratio > 1 indicates a positive association and an odds ratio < 1 indicates a negative association. Statistical significance was determined by an adjusted p-value ≤ 0.05 using Fisher’s exact test with Benjamini-Hochberg p-value adjustment. *See also Table S1*. **(E)** Genomic tracks showing examples of persistent DMR hypermethylation located in CpG islands and promoters. **(F)** Z-score normalization of DNA methylation levels in wild-type, *Dnmt1^tet/tet^*, doxycycline-treated *Dnmt1^tet/tet^* mESCs and *Dnmt1^tet/tet^* mESCs after removing doxycycline (21-day recovery period; D1^t/t^ Rec) for *Dnmt1^tet/tet^* hypermethylated DMRs that showed < 10% increase in DNA methylation in doxycycline-treated *Dnmt1^tet/tet^* versus wild-type mESCs in **B** i.e. non-persistent DMR hypermethylation. **(G)** Genomic tracks showing DMRs in **F** that regained a hypermethylated state (≥ 10% increase versus WT; first two panels) or remained hypomethylated (< 10% increase versus WT; third panel) following doxycycline treatment recovery.

We next asked whether the non-persistent hypermethylated DMRs—those showing less than 10% methylation increase in dox-treated *Dnmt1^tet/tet^* mESCs relative to wild-type (Figure 5B)—would regain their hypermethylated state upon reinstating DNMT1 overexpression. To address this, doxycycline was removed from *Dnmt1^tet/tet^* mESCs and cells were allowed to recover for 21 days before profiling DNA methylation genome-wide^33^. Although DNMT1 is re-expressed to its original level within ∼2 days of doxycycline withdrawal^31^, the 21-day recovery period was chosen to allow DNA methylation profiles to stabilize following DNMT1 re-expression^33^. Among the non-persistent DMRs, the majority regained hypermethylation following the recovery period, while a minority remained hypomethylated (Figures 5F–G). This indicates that DNMT1 overexpression is largely sufficient to re-establish hypermethylation at these loci once reintroduced, consistent with DNMT1 being the primary driver of hypermethylation at these sites. The subset of DMRs that remained hypomethylated after doxycycline recovery may reflect loci where the local chromatin or sequence context prevents efficient DNMT1-mediated re-methylation, or where a longer recovery period would be required.

Altogether, these findings highlight potential limitations of DNMT1-specific inhibition in cancer therapy, which are typically administered on cyclical, temporary schedules. While we show that hypermethylation at a subset of loci is effectively erased by DNMT1 depletion and is not re-established upon reinstating DNMT1 overexpression, the majority of hypermethylation either persists despite DNMT1 loss or is restored once DNMT1 is overexpressed again. Complete and durable erasure of hypermethylation at persistent loci may therefore require pan-DNMT inhibitors, and extended treatment regimens may be needed to achieve sufficient cytotoxicity to prevent re-emergence of hypermethylation.

### Conserved promoter hypermethylation between *Dnmt1^tet/tet^* mESCs and human cancer

*Dnmt1^tet/tet^* hypermethylated promoter DMRs were enriched for cancer-associated genes (Figure 3A), consistent with DNMT1 overexpression being associated with promoter hypermethylation in human cancer^24,42–44^. To further evaluate the translational relevance of these findings, we asked whether human cancer samples would show promoter hypermethylation in the same regions identified in *Dnmt1^tet/tet^* mESCs. Accordingly, we retrieved publicly available whole-genome DNA methylation datasets from both male and female human samples, which included gastric cancer (pre-cancerous intestinal metaplasia and dysplasia, and cancerous adenocarcinoma)^97^, prostate cancer (LNCaP^98^ and PC3^99^ cell lines, and prostate adenocarcinoma^100^), hepatocellular carcinoma^101^, renal cell carcinoma^102^, pancreatic ductal adenocarcinoma^103^, breast cancer (MCF-7^99^ and HCC1954^104^ cell lines), cervical cancer^105^ and glioblastoma^106^, each with matched normal controls (Data S7). Differential DNA methylation between cancer and normal samples was then quantified in human genomic regions corresponding to the *Dnmt1^tet/tet^* hypermethylated promoter DMRs, defining changes of at least 10% as biologically meaningful and including only regions with at least five mapped CpGs. This analysis revealed that although most *Dnmt1^tet/tet^* hypermethylated promoter DMRs were unchanged in the human cancer samples, many of them were hypermethylated (median frequency of 14%), while relatively few of them were hypomethylated (median frequency of 0.3%) (Figures 6A, 7A; Table S2). In other words, if these regions were altered in the human cancer samples, they generally did so in the same direction as in *Dnmt1^tet/tet^* mESCs. The highest frequencies of conserved *Dnmt1^tet/tet^* promoter hypermethylation were observed in gastric intestinal metaplasia (14%), dysplasia (14%) and adenocarcinoma (17%), the prostate cancer cell line PC3 (21%), and the breast cancer cell lines MCF-7 (24%) and HCC1954 (15%) (Figures 6A, 7A; Table S2). To determine whether the evolutionary conservation of these promoter DMRs might influence their propensity to undergo hypermethylation in both *Dnmt1^tet/tet^* mESCs and human cancers, we next measured their sequence homology between mouse and human. Because promoters are generally well conserved to begin with, we did not anticipate major differences in sequence identity between *Dnmt1^tet/tet^* hypermethylated promoter DMRs that were or were not conserved in human cancers. Nonetheless, regions that showed hypermethylation in at least one cancer dataset exhibited modest but statistically significant increases in sequence homology (mean = 60.0%) compared with those that were not hypermethylated in any cancer dataset (mean = 57.5%) and compared with all mapped regions (mean = 58.5%) (Figure 7B). These findings suggest that even within this already conserved promoter space, higher sequence homology may slightly increase the likelihood of concordant promoter hypermethylation between *Dnmt1^tet/tet^* mESCs and human cancers.

**Figure 6.**
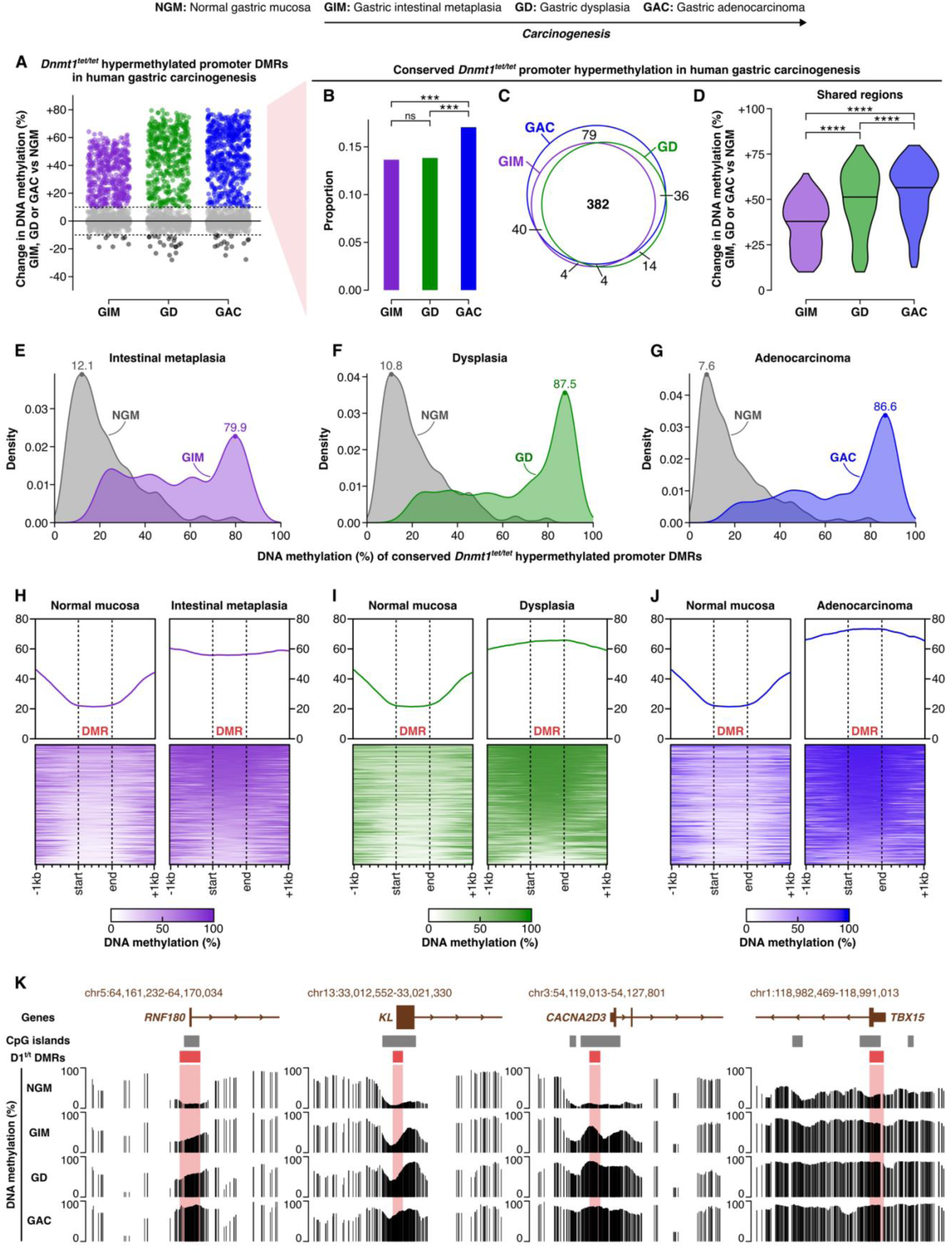
Conserved *Dnmt1^tet/tet^* promoter hypermethylation throughout human gastric carcinogenesis. **(A)** Change in DNA methylation in human male gastric intestinal metaplasia (GIM; n=3), dysplasia (GD; n=2) and adenocarcinoma (GAC; n=1, male) versus matched normal mucosa (NGM; n=3) for human genomic regions corresponding to the *Dnmt1^tet/tet^* hypermethylated promoter DMRs. GIM, GD, GAC and NGM datasets were obtained via the UCSC table browser. *See also Data S7*. **(B)** Proportion of regions in **A** that showed ≥ 10% increase in DNA methylation in GIM, GD or GAC versus NGM i.e. conserved *Dnmt1^tet/tet^* promoter hypermethylation. Statistical comparison was conducted using Pearson’s chi-squared test with Benjamini-Hochberg p-value adjustment. *** p-value ≤ 0.001, ns: not significant. *See also Table S2*. **(C)** Number of regions of conserved *Dnmt1^tet/tet^* promoter hypermethylation identified in **B** that were specific or shared among GIM, GD and GAC. **(D)** Change in DNA methylation in GIM, GD and GAC versus NGM for the shared regions of conserved *Dnmt1^tet/tet^* promoter hypermethylation identified in **C**. Statistical comparisons were conducted using the pairwise Wilcoxon test with Benjamini-Hochberg p-value adjustment. **** p-value ≤ 0.0001. **(E–G)** DNA methylation levels in NGM and GIM, GD or GAC for regions of conserved *Dnmt1^tet/tet^* promoter hypermethylation in GIM, GD or GAC, respectively. The dot on each density plot indicates the methylation level at the distribution’s highest peak (i.e. primary mode). **(H)** DNA methylation levels in NGM and GIM for regions of conserved *Dnmt1^tet/tet^* promoter hypermethylation in GIM and their flanking regions (± 1kb). **(I–J)** Same as **H** but for GD or GAC, respectively. **(K)** Genomic tracks showing examples of shared regions of conserved *Dnmt1^tet/tet^* promoter hypermethylation in GIM, GD and GAC that were identified in **C**.

**Figure 7.**
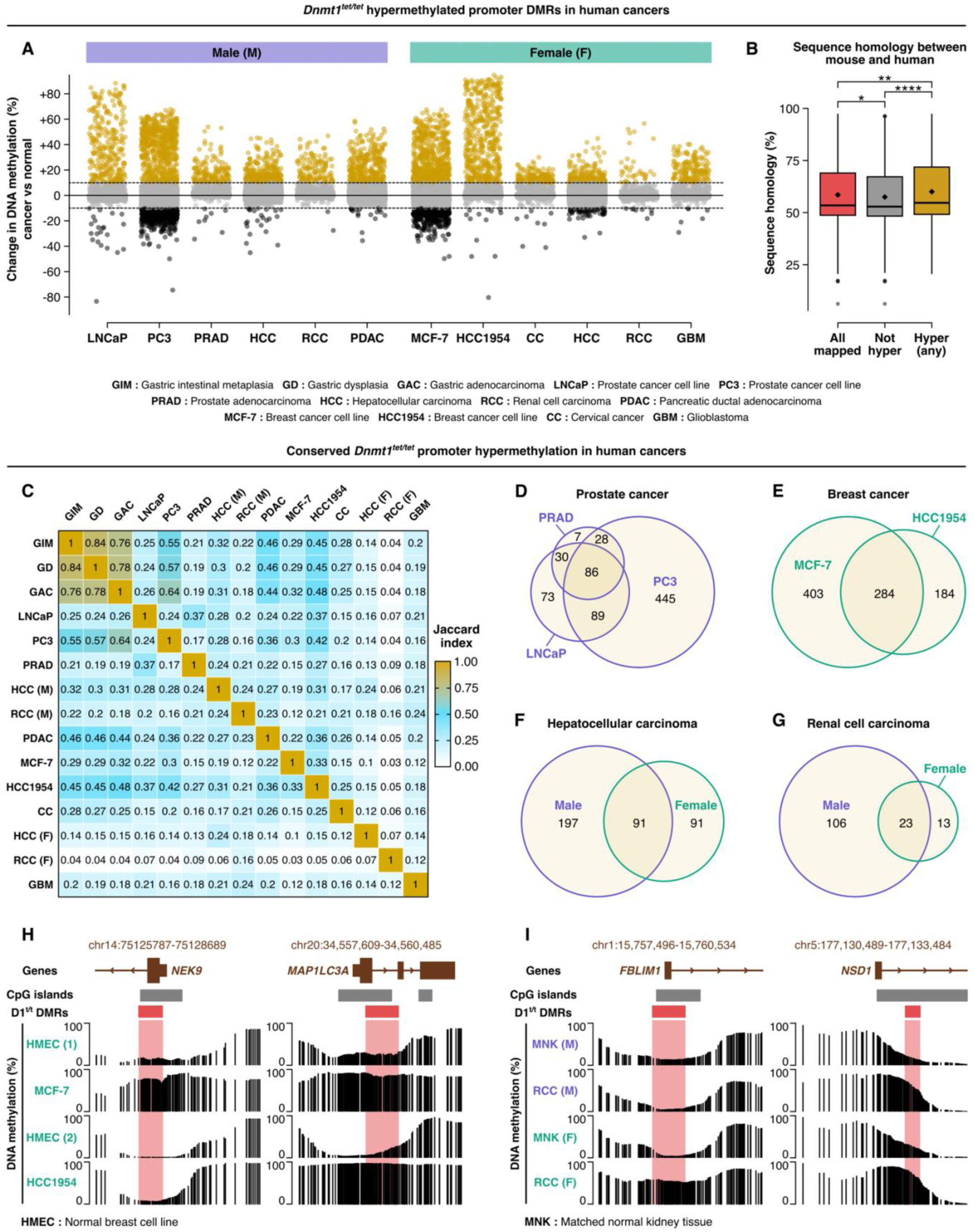
Conserved promoter hypermethylation between *Dnmt1^tet/tet^* mESCs and human cancer. **(A)** Change in DNA methylation in human cancer versus matched normal samples for human genomic regions corresponding to the *Dnmt1^tet/tet^* hypermethylated promoter DMRs. Datasets were obtained via the UCSC table browser. *See also Data S7*. **(B)** Sequence homology percentage between mouse and human for *Dnmt1^tet/tet^* hypermethylated promoter DMRs for all that were mapped in the human DNA methylation datasets, for ones that were not hypermethylated in any human cancer dataset and for ones that were hypermethylated in at least one human cancer dataset. Statistical comparisons were conducted using the pairwise Wilcoxon test with Benjamini-Hochberg p-value adjustment. * p-value ≤ 0.05, ** p-value ≤ 0.01, **** p-value ≤ 0.0001. **(C)** Pairwise Jaccard indices quantifying cross-cancer similarity of conserved *Dnmt1^tet/tet^* promoter hypermethylation i.e. regions in **A** and Figure 6A that showed ≥ 10% increase in DNA methylation. An index of 1 signifies identical sets of regions and an index of 0 signifies unique sets of regions. *See also Tables S2 and S3*. **(D)** Number of regions of conserved *Dnmt1^tet/tet^* promoter hypermethylation that were specific or shared among prostate cancer samples (LNCaP, PC3, PRAD). **(E–G)** Same as **D** but for breast cancer samples (MCF-7, HCC1954), HCC samples (male, female) or RCC samples (male, female), respectively. Regions on chromosomes X and Y were not included in the analyses in **C**, **F** and **G**. **(H)** Genomic tracks showing a specific and shared region of conserved *Dnmt1^tet/tet^* promoter hypermethylation among breast cancer samples (MCF-7, HCC1954). **(I)** Same as **H** but for male and female RCC.

The human gastric cancer datasets, which uniquely included both pre-cancerous and cancerous conditions, enabled the analysis of conserved *Dnmt1^tet/tet^* promoter hypermethylation throughout carcinogenesis progression, from intestinal metaplasia to dysplasia to adenocarcinoma^97^. Conserved *Dnmt1^tet/tet^* promoter hypermethylation was more abundant in adenocarcinoma than in intestinal metaplasia and dysplasia (Figure 6B), with 537, 430 and 436 regions, respectively (Table S2). Remarkably, the majority of hypermethylated regions in adenocarcinoma (n = 382) were also hypermethylated in both pre-cancerous conditions (Figure 6C) and exhibited a stepwise increase in hypermethylation throughout carcinogenesis progression (Figure 6D). This early initiation and progressive intensification of promoter hypermethylation could be indicative of epigenetic priming^107^, supporting the predictive power of epigenetic biomarkers in early disease detection^27^. To address the possibility that hypermethylation in cancer versus normal samples at these regions reflect shifts in cellular composition rather than genuine cancer-associated hypermethylation, we examined the distribution of methylation levels across conserved *Dnmt1^tet/tet^* promoter hypermethylated regions in matched normal gastric mucosa (NGM) and each cancer condition (GIM, GD or GAC). The resulting density plots revealed distinctly polarized distributions, with primary modes at 12.1% and 79.9% for NGM and GIM (Figure 6E), 10.8% and 87.5% for NGM and GD (Figure 6F), and 7.6% and 86.6% for NGM and GAC (Figure 6G), respectively. The prominence of these peaks indicate generally low methylation heterogeneity at these regions across both normal and cancer samples, arguing against cellular composition shifts as a major confounding factor and instead supporting a consistent gain of methylation within the cancer cell population. Moreover, hypermethylation in gastric intestinal metaplasia, dysplasia and adenocarcinoma extended beyond the regional boundaries defined in *Dnmt1^tet/tet^* mESCs yet was most pronounced within those boundaries (Figure 6H–J), suggesting that *Dnmt1^tet/tet^* mESCs captured core epigenetic signatures nested within broader domains of dysregulation. Promoters of *RNF180*^108^, *KL*^109^, *CACNA2D3*^110^ and *TBX15*^93^, genes implicated in cancer, exemplified these dynamics. *RNF180* displayed hypermethylation mostly confined to the region identified in *Dnmt1^tet/tet^* mESCs, whereas *KL*, *CACNA2D3* and *TBX15* exhibited much broader hypermethylation, with *RNF180*, *KL* and *CACNA2D3* showing a progressive increase in hypermethylation across stages of carcinogenesis (Figure 6K).

Next, to ask whether the regions conserved between our model and each cancer type are shared across cancers or specific to particular cancer types, we assessed the cross-cancer specificity of conserved *Dnmt1^tet/tet^* promoter hypermethylation across human cancer samples by computing pairwise Jaccard indices^111^. A Jaccard index quantifies the similarity between two sets of elements, with an index of 1 indicating identical sets and an index of 0 indicating unique sets. Chromosomes X and Y were excluded from this analysis to allow comparisons between male and female samples. Interestingly, conserved *Dnmt1^tet/tet^* promoter hypermethylation mostly showed limited cross-cancer overlap as nearly all pairwise comparisons yielded a Jaccard index below 0.5, with a median of 0.21 overall (Figure 7C; Table S3). Jaccard indices above 0.5 were only observed for comparisons involving gastric intestinal metaplasia, dysplasia and adenocarcinoma; they expectedly had strong indices with one another, ranging from 0.76 to 0.84, but also had moderately high indices of 0.55, 0.57 and 0.64, respectively, with the prostate cancer cell line PC3 (Figure 7C; Table S3). Even cancer samples from the same tissue of origin (prostate: LNCaP, PC3, PRAD; breast: MCF-7, HCC1954), or male and female samples of the same type of cancer (HCC, RCC), shared relatively low Jaccard indices that ranged from 0.16 to 0.37 (Figure 7C; Table S3). Many *Dnmt1^tet/tet^* hypermethylated promoter DMRs were conserved in only one of LNCaP, PC3, or PRAD (Figure 7D); MCF-7 or HCC1954 (Figure 7E); male or female HCC (Figure 7F); and male or female RCC (Figure 7G). For example, we observed *NEK9* promoter hypermethylation in the breast cancer cell line MCF-7 but not in HCC1954, and *FBLIM1* promoter hypermethylation in female RCC but not in male RCC, whereas normal samples showed similar methylation profiles (Figure 7H–I). Together, these findings illustrate how DNA methylation can discriminate distinct^112^ and related types of cancer^113^ with sex specificity^114^. However, conserved *Dnmt1^tet/tet^* promoter hypermethylation in human cancer samples also showed points of commonality, reflecting how cancers can share hallmark mechanisms^24^. For instance, the *MAP1LC3A* promoter was hypermethylated in both breast cancer cell lines, and the *NSD1* promoter was hypermethylated in both male and female RCC (Figure 7H–I). Moreover, the *TBX15* promoter, which showed conserved *Dnmt1^tet/tet^* hypermethylation in gastric intestinal metaplasia, dysplasia and adenocarcinoma (Figure 6K), was also hypermethylated in LNCaP, PC3, PRAD, male HCC, PDAC, MCF-7, HCC1954 and CC (Figure S2).

While DNMT1 overexpression has been documented in many of the cancer types we examined^24,42–44,115,116^, it was not assessed in the specific studies from which we obtained the DNA methylation datasets, with the exception of glioblastoma, where Raiber et al. reported DNMT1 polyploidy indicative of overexpression^106^. Although DNMT1 overexpression is a widely recognized feature of many human cancers, precise fold-change measurements are infrequently reported, as many studies rely on immunoreactivity-based assessments rather than quantitative RNA or protein analyses. Nonetheless, breast cancer studies by Mirza et al. and Agoston et al. provide illustrative examples of the magnitude and regulatory mechanisms of DNMT1 dysregulation in human cancer: Mirza et al. reported 1.2- to 4.4-fold increases in DNMT1 mRNA in invasive ductal carcinoma tumors relative to matched normal tissues^115^, and Agoston et al. observed a ∼2-fold increase in DNMT1 mRNA in the MCF-7 cell line relative to normal human mammary epithelial cells (HMECs)^116^. Despite the moderate transcriptional increase, Agoston et al. demonstrated that DNMT1 protein was elevated ∼6-fold in MCF-7 cells, which was attributable to impaired proteasome-mediated degradation^116^—illustrating that post-translational stabilization can substantially amplify DNMT1 protein accumulation in cancer beyond what mRNA levels alone would predict. By contrast, the ∼7-fold increase in DNMT1 mRNA in *Dnmt1^tet/tet^* mESCs yielded only a ∼3-fold protein elevation, which suggests that post-translational turnover remains efficient in these otherwise healthy cells and that DNMT1 protein levels in our model represent a conservative estimate relative to what may accumulate in cancer through combined transcriptional and post-translational dysregulation. Inclusion of human cancer studies in our analyses was primarily guided by the availability of high-quality DNA methylation datasets with clear sample annotations rather than documented DNMT1 overexpression. Despite this caveat, the ability of *Dnmt1^tet/tet^* mESCs to partially recapitulate cancer-associated promoter hypermethylation at a meaningful subset of loci underscores their translational value, while the marked variability in overlap across cancer types and sexes highlights the specificity and diagnostic potential of DNA methylation-based biomarkers. A key challenge moving forward will be to distinguish DNA methylation alterations that act as causal drivers from those that represent downstream footprints of disease processes.

To conclude, we employed *Dnmt1^tet/tet^* mESCs to study the effect of DNMT1 overexpression on DNA methylation regulation with high resolution in a controlled biological setting. Compared to wild-type mESCs, *Dnmt1^tet/tet^* mESCs exhibited global hypomethylation, localized hypermethylation and increased CpG-to-CpG variability—features shared with cancer methylomes^25,26^—as well as reduced DNMT3A and DNMT3B levels. These findings suggest that aberrant DNMT1 dosage influences DNA methylation not only through its own catalytic activity but also by altering the balance among DNMTs and the fidelity of maintenance processes. DNA methylation alterations in promoters were linked to developmental, neuronal, cardiac and cancer-associated pathways, correlated with differential gene expression and intersected binding sites of key transcriptional regulators (HDAC1, E2F1, CREBBP, MBD1, KMT2C–D, PRC2), indicating that DNMT1 overexpression may reshape regulatory networks through multiple, context-dependent mechanistic layers. Interestingly, a subset of hypermethylated regions persisted following DNMT1 depletion, raising the possibility that once established, some hypermethylation may be maintained by residual DNMT3A and DNMT3B activity or stabilized by other chromatin features. Future studies dissecting the temporal sequence of events, the contributions of individual DNMTs, and the interplay with transcriptional regulators and chromatin states will be essential for clarifying the underlying mechanisms of DNMT1 overexpression. Importantly, however, we show that DNMT1 depletion erased the majority of hypermethylation, which was then largely regained once DNMT1 overexpression was reinstated, supporting DNMT1’s central role in driving hypermethylation events in *Dnmt1^tet/tet^* mESCs. Moreover, we found that various human cancer samples showed hypermethylation at a substantial portion of the hypermethylated promoter regions identified in *Dnmt1^tet/tet^* mESCs, suggesting that at least some loci affected in this model may be relevant in human cancers that overexpress DNMT1^24,42–44^. While our focus was on cancer, the gene pathways enriched among differentially methylated promoters in *Dnmt1^tet/tet^* mESCs point to potential relevance in other human diseases associated with DNMT1 overexpression, such as schizophrenia^54^, epilepsy^55^, bipolar disorder^56^ and cardiomyopathy^53^. Extending this research into more complex developmental and disease systems will help expand its translational impact. Overall, our work positions *Dnmt1^tet/tet^* mESCs as a valuable model for examining how excess DNMT1 perturbs DNA methylation regulation and contributes to pathological processes, highlighting specific genomic regions and candidate mechanisms for deeper exploration.

### Limitations

Some limitations should be acknowledged when interpreting the findings of this study. First, experiments were only performed in mESCs, a highly plastic cellular context that may respond differently to DNMT1 overexpression than somatic or diseased cells and tissues. Second, EM-seq does not distinguish between 5-methylcytosine and 5-hydroxymethylcytosine; however, hydroxymethylation is unlikely to substantially influence our conclusions given its lack of maintenance across cell divisions^117,118^, the repeated passaging of mESCs prior to EM-seq and the unchanged expression of *Tet1*, *Tet2* and *Tet3*. Third, while the proposed relationships linking DNA methylation alterations with reduced DNMT3 levels or with enrichment of transcriptional regulator binding sites are evidence-based and plausible, they remain correlative and will require further testing to assess mechanistic causality. Fourth, comparisons with human cancer datasets revealed compelling parallels with *Dnmt1^tet/tet^* mESCs but do not establish functional equivalence given biological and technical differences. Fifth, the use of male *Dnmt1^tet/tet^* mESCs precluded sex-specific analyses, although our human cancer analyses incorporated both male and female samples. Finally, DNMT1 overexpression was studied in mESCs using only one experimental system, therefore, we cannot exclude the possibility that some effects may be model-specific. However, our findings are consistent with established DNMT1 biology and with observations described in other biological contexts, suggesting that our core conclusions are unlikely to be unique to this model. Despite these limitations, the depth and consistency of the DNA methylation alterations we observed, along with their convergence across independent human cancer methylomes, support the robustness of our conclusions and provide a strong foundation for targeted mechanistic and translational follow-up studies.

## Material and Methods

### Culture of mESCs and doxycycline treatment

Cell culture of wild-type (R1) and *Dnmt1*^tet/tet^ mESCs in serum-containing medium, and doxycycline treatment and recovery of *Dnmt1*^tet/tet^ mESCs (i.e. 2 ug/mL for 6 days) were performed as described in Elder et al.^33^

### Enzymatic methyl sequencing

DNA extraction and EM-seq of wild-type mESCs were performed in triplicates as described in Elder et al.^33^ We obtained 514M, 415M and 473M raw reads per sample. Raw reads were processed using the GenPipes^119^ DNA methylation sequencing pipeline, also described in Elder et al.^33^, to obtain CpG coverage and methylation calls. EM-seq datasets (in triplicates) for *Dnmt1*^tet/tet^ and doxycycline-treated mESCs were repurposed from Elder et al.^33^ for this study. Overall, an average of 15,714,082 CpGs were covered per sample (∼75% of genomic CpGs), with ∼86% sequenced at ≥ 10X, reflecting high coverage and sequencing depth. The DML.test function from the DSS R package v2.58.0^41^ was then used to produce smoothed DNA methylation levels for each experimental condition, normalized for coverage and biological variability across replicates. Smoothing span was set at 500 bp (default). Because smoothed DNA methylation levels needed to be comparable across all three experimental conditions, the DML.test function and its internal helper function getBSseqIndex were adapted to include a third group during the CpG filtering steps (original functions only accept two groups). This ensured that the same CpGs were retained in all three groups. The adapted functions can be found here: https://github.com/elderelizabeth/misc-methyl.git.

### Differentially methylated regions

Differentially methylated regions (DMRs) in *Dnmt1*^tet/tet^ versus wild-type mESCs were identified using the DSS R package v2.58.0^41^. All steps in the DSS workflow were followed as instructed. DMR identification parameters were set at ≥ 100 bp for length, ≥ 5 for the number of CpGs, ≥ 50% for the proportion of CpGs showing significant (adjusted p-value ≤ 0.01) change ≥ 10%, and ≤ 100 bp for the merging distance. Then, DMRs were filtered to only keep those that showed a regional mean change ≥ 10%, with an increase ≥ 10% defining hypermethylated DMRs and a decrease ≥ 10% defining hypomethylated DMRs. To measure CpG density, we used all genomic CpGs from the mm10 reference genome, not only the CpGs that were sequenced. Genomic CpGs were retrieved from the BSgenome.Mmusculus.UCSC.mm10 R package v1.4.3. To measure the distance between DMRs and the nearest transcription start site (TSS), genomic coordinates of TSSs were obtained from the TxDb.Mmusculus.UCSC.mm10.knownGene R package v3.10.0, only retaining those that had an Entrez ID. To identify persistent or non-persistent DMR hypermethylation upon DNMT1 depletion in *Dnmt1*^tet/tet^ mESCs, we computed regional mean DNA methylation levels of *Dnmt1*^tet/tet^ hypermethylated DMRs in doxycycline-treated *Dnmt1*^tet/tet^ mESCs. Those that showed a sustained increase ≥ 10% in doxycycline-treated *Dnmt1*^tet/tet^ versus wild-type mESCs were defined as being persistently hypermethylated, while the rest were classified as non-persistent. To assess whether non-persistent DMRs could regain hypermethylation upon reinstating DNMT1 overexpression, we computed regional mean DNA methylation levels of these DMRs in *Dnmt1^tet/tet^* mESCs following a 21-day doxycycline withdrawal recovery period. Non-persistent DMRs showing a methylation increase ≥ 10% relative to wild-type mESCs after recovery were classified as having regained hypermethylation, while the rest were considered to have undergone stable hypomethylation.

### Genomic feature annotations

The mouse mm10 reference genome was used for all genomic feature annotations. Promoter and gene body regions were obtained using the GenomicFeatures v1.62.0 and TxDb.Mmusculus.UCSC.mm10.knownGene v3.10.0 R packages, only retaining those that had an Entrez ID. We defined promoters as starting 1000 bp upstream and ending 500 bp downstream of TSS. We then divided promoter regions into core (±50 bp from TSS), proximal (±50 bp to ±500 bp from TSS) and distal (-500 bp to -1000 bp from TSS) subregions. Intergenic regions were defined as all genomic regions not covered by a promoter or gene body. CpG islands, CpG island shores, CpG island shelves and open sea regions (annotatr R package v1.36.0), imprinted regions (n=26 curated from literature), retrotransposons (LINEs, SINEs and LTRs from RepeatMasker with a Smith-Waterman score ≥ 1000) were defined as described in Elder et al.^33^

### mRNA sequencing

RNA extraction and messenger RNA sequencing (mRNA-seq) of wild-type mESCs were performed in triplicates as described in Elder et al.^33^ We obtained 29M, 28M and 28M raw reads per sample. Raw reads were processed using the GenPipes^119^ RNA sequencing pipeline, also described in Elder et al.^33^, to obtain raw counts. mRNA-seq datasets (in triplicates) for *Dnmt1*^tet/tet^ and doxycycline-treated mESCs were repurposed from Elder et al.^33^ for this study.

### Differentially expressed genes

Since we aimed to correlate gene expression with promoter DNA methylation, raw counts were filtered to only retain genes with available promoter DNA methylation levels in our EM-seq datasets. See the “Genomic feature annotations” section above for how we retrieved and defined promoter regions. Then, the DESeq2 R package v1.50.2^120^ was used to compute normalized counts and identify differentially expressed genes (DEGs). All steps in the DESeq2 workflow were followed as instructed, including the facultative steps of filtering out low raw counts (i.e. only genes with ≥ 10 counts in ≥ 3 samples were kept) and log_2_ fold change shrinking (type = “apeglm”). DEGs were retained if their adjusted p-value was ≤ 0.05 and log_2_ fold change was either ≥ 1 (upregulated) or ≤ -1 (downregulated).

### Western blots

Protein extraction and western blots were performed as described in Elder et al.^33^, except that here we captured images of Ponceau S-stained membranes. Membranes were incubated with Ponceau S (Thermo Fisher Scientific, A40000279) for 1 minute, then rinsed with water to remove excess Ponceau S and imaged using the ChemiDoc system. Membranes were subsequently washed in Tris-buffered saline containing 0.1% Tween-20 until the Ponceau S staining had fully disappeared, after which the western blot procedure was resumed. Quantification of western blot band intensities was conducted using ImageJ software by measuring integrated densities, which were corrected with background signals and normalized to total protein signals from the Ponceau S staining images. See Data S2 for detailed quantification calculations.

<u>Primary antibodies:</u> anti-DNMT1 (1/1000 dilution; Cell Signaling Technology, 5032), anti-DNMT3A (1/1000 dilution; Cell Signaling Technology, 49768), anti-DNMT3B (1/1000 dilution; Abcam, ab2851) and anti-ACTB (beta-actin; 1/1000 dilution; Cell Signaling Technology, 4970)

<u>Secondary antibody:</u> anti-rabbit IgG HRP-linked (1/10000 dilution; Cell Signaling Technology, 7074)

### Calculation of posterior mean of alpha values

Posterior mean of alpha values of genome-wide bins of 101 CpGs were calculated using the plotAlphaDistributionOneChr function from the MethylSeekR R package v1.50.0^48^. Since the original plotAlphaDistributionOneChr function can only process one chromosome at a time and only outputs a histogram figure, we adapted the function to process all chromosomes sequentially and to output the numeric values in addition to figures. The adapted function can be found here: https://github.com/elderelizabeth/misc-methyl.git.

### Enrichment of KEGG pathways

Enrichment of KEGG pathways in genes associated with *Dnmt1*^tet/tet^ promoter DMRs was conducted using the enrichKEGG function from the clusterProfiler R package v4.18.2. The “minGSSize” parameter was set at 10, the “maxGSSize” parameter was set at 1000 and the “universe” was defined as all genes with available promoter DNA methylation levels. Enriched pathways with a BH-adjusted p-value ≤ 0.05 were retained.

### Enrichment of transcriptional regulator binding sites

Enrichment of transcriptional regulator binding sites overlapping with *Dnmt1*^tet/tet^ promoter DMRs was conducted using the enrichment function from the ReMapEnrich R package v0.99.0. The “shuffles” parameter was set at 5000, “fractionQuery” at 0.1 and “fractionCatalog” at 0.1. The universe was defined as all promoter regions with available promoter DNA methylation levels. The query was defined as either the *Dnmt1*^tet/tet^ hypermethylated or hypomethylated promoter DMRs, which were reduced to fit within the universe using the redefineUserSets function from the LOLA R package v1.40.0. The catalog was defined as all ChIP-seq (chromatin immunoprecipitation sequencing) datasets of transcriptional regulator binding sites, which were obtained from the mouse ReMap 2022 catalog^72^. Enriched datasets with a BH-adjusted p-value ≤ 0.05 were retained.

### Analysis of human DNA methylation datasets

The human (GRCh38/hg38) whole-genome DNA methylation datasets (i.e. CpG coverage and methylation calls) were obtained through the UCSC table browser; sample annotations and access links are provided in Data S7. These datasets were originally produced in various studies^97–106^ but were processed by the Smith Lab^121^. The DML.test function from the DSS R package v2.58.0^41^ was then used to produce smoothed DNA methylation levels as described in the “Enzymatic methyl sequencing” section above. The original versions of DML.test and its internal helper function getBSseqIndex were used for datasets with two groups (i.e. normal, cancer). For gastric cancer datasets, which contained four groups (i.e. normal mucosa, intestinal metaplasia, dysplasia, adenocarcinoma), DML.test and getBSseqIndex were adapted to accommodate four groups during the CpG filtering steps, ensuring that the same CpGs were retained across all groups and that smoothed DNA methylation levels were comparable. The adapted functions can be found here: https://github.com/elderelizabeth/misc-methyl.git. To evaluate conserved *Dnmt1*^tet/tet^ promoter hypermethylation in human cancer, human (GRCh38/hg38) genomic coordinates corresponding to the *Dnmt1*^tet/tet^ hypermethylated DMRs were obtained using the UCSC LiftOver tool. *Dnmt1*^tet/tet^ promoter hypermethylation was considered conserved in human cancer if the corresponding human genomic regions contained ≥ 5 mapped CpGs in the human DNA methylation datasets and exhibited a ≥ 10% increase in DNA methylation in cancer relative to matched normal samples.

### Sequence homology analysis

Hypermethylated promoter DMRs identified in *Dnmt1^tet/tet^* mESCs (mm10) were lifted over to human genome coordinates (hg38) using the UCSC LiftOver tool, and DMRs that failed liftover were excluded. DNA sequences underlying each DMR were extracted from the mm10 and hg38 reference genomes using the BSgenome R packages. Pairwise global sequence alignments between orthologous mouse and human DMR sequences were performed using pwalign::pairwiseAlignment, and percent sequence identity was computed for each pair. Then, sequence homology percentages for all DMRs that were mapped in the human DNA methylation datasets, for DMRs that were not hypermethylated in any cancer dataset, and for DMRs that were hypermethylated in at least one cancer dataset were compared using the pairwise Wilcoxon test with Benjamini-Hochberg p-value adjustment.

### Pairwise Jaccard similarity analysis

Pairwise Jaccard similarity analysis of genomic regions showing conserved *Dnmt1*^tet/tet^ promoter hypermethylation across different human cancers was conducted in R v4.5.2. The Jaccard similarity index^111^ was computed as:

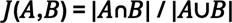

***J*** is the Jaccard similarity index, ***A*** and ***B*** are the two sets of regions being compared, **|*A*∩*B*|** is the count of shared regions, and **|*A*∪*B*|** is the total count of unique regions in both sets. A pairwise matrix of indices was then constructed by computing

***J*(*A*,*B*)** for all combinations of ***A*** and ***B***. Regions on chromosomes X and Y were removed from all sets to enable similarity analysis between male and female samples and to ensure that overall indices were comparable.

### Statistical analyses

Statistical analyses were conducted in R v4.5.2 using functions from the stats package v4.5.2, except where otherwise specified. The Shapiro-Wilk test for normality (shapiro.test function; performed on residuals generated by the aov and residuals functions) and Levene’s test for homogeneity of variance (leveneTest function, car package v3.1.3) were used to decide whether parametric or non-parametric statistical tests were appropriate. Parametric testing was only conducted when comparing three groups of data, for which we used one-way ANOVA (aov function) followed by Tukey’s HSD test (TukeyHSD function). For non-parametric testing, the Wilcoxon rank sum test (wilcox.test function) was used when comparing two groups of data, and the Kruskal-Wallis test (kruskal.test function) followed by pairwise Wilcoxon rank sum tests (pairwise.wilcox.test function) were used when comparing three groups of data. For comparing categorical data, the chisq.test function was used for Pearson’s chi-squared tests and the fisher.test function was used for computing odds ratios and corresponding p-values. When appropriate, Benjamini-Hochberg p-value adjustment for multiple testing was applied either within the statistical test function if possible or subsequently using the p.adjust function. Spearman’s correlation was conducted using the stat_cor function from the ggpubr package v0.6.2. Dixon’s Q test was employed for detecting outliers using the dixon.test function from the outliers package v0.15.

## Acknowledgements

We thank the members of the McGraw laboratory for their feedback and support, Génome Québec for sequencing services, as well as Gregor Andelfinger (University of Montreal) and J. Richard Chaillet (University of Pittsburgh) for gifting R1 and *Dnmt1^tet/tet^* mESCs, respectively. We are also thankful to the RQR (Réseau Québécois en reproduction), the CRRF (Centre de recherche en reproduction et fertilité), the ODISÉ (Origines développementales et intergénérationnelles de la santé des enfants) network, and ThéCell (Réseau de thérapie cellulaire) for supporting student training and this research. The graphical abstract was created in BioRender.com.

## Funding

This work was supported by a Natural Sciences and Engineering Research Council of Canada grant to S.M. (RGPIN-2023-04559) and bursary to E.E., and a Fonds de recherche du Québec senior salary award to S.M. and bursary to E.E.

## Author contributions

Conceptualization & methodology: E.E., S.M.; Investigation: E.E.; Formal analysis: E.E., A.L.; Writing–original draft: E.E., S.M.; Writing–review & editing: E.E., A.L., S.M.; Visualization: E.E.; Supervision: S.M.; Funding acquisition: S.M.

## Competing interests

None declared.

## Data and materials availability

EM-seq and mRNA-seq datasets for R1 mESCs (wild type) are deposited in the Gene Expression Omnibus (GEO) under the accession numbers GSE306782 and GSE306781, and those for *Dnmt1^tet/tet^* mESCs (doxycycline-treated, after doxycycline recovery) under GSE267053. Binding sites of transcriptional regulators can be retrieved from the mouse ReMap 2022 dataset catalog^72^. Human DNA methylation datasets were obtained via the UCSC table browser; access links are listed in Data S7. We do not report original code but our adapted functions from the R packages DSS (getBSseqIndex, DMLtest) and MethylSeekR (plotAlphaDistributionOneChr) are available here: https://github.com/elderelizabeth/misc-methyl.git. Scripts for data analysis can be provided upon request to lead contact. All materials used in this study are commercially available or can be provided upon request to lead contact.

## Supplementary Material

Supplementary material that support this manuscript include two figures, three tables and seven data.

## Notes

### Competing Interest Statement

The authors have declared no competing interest.

### Summary of Updates

The title and abstract were improved. No other changes were made.

