## Supplementary materials for "DNMT1 overexpression disrupts DNA methylation homeostasis in mouse embryonic stem cells"

**This PDF file includes:**

- Figures S1 to S2
- Tables S1 to S3

**Other supplementary materials include:**

- Data S1 to S7

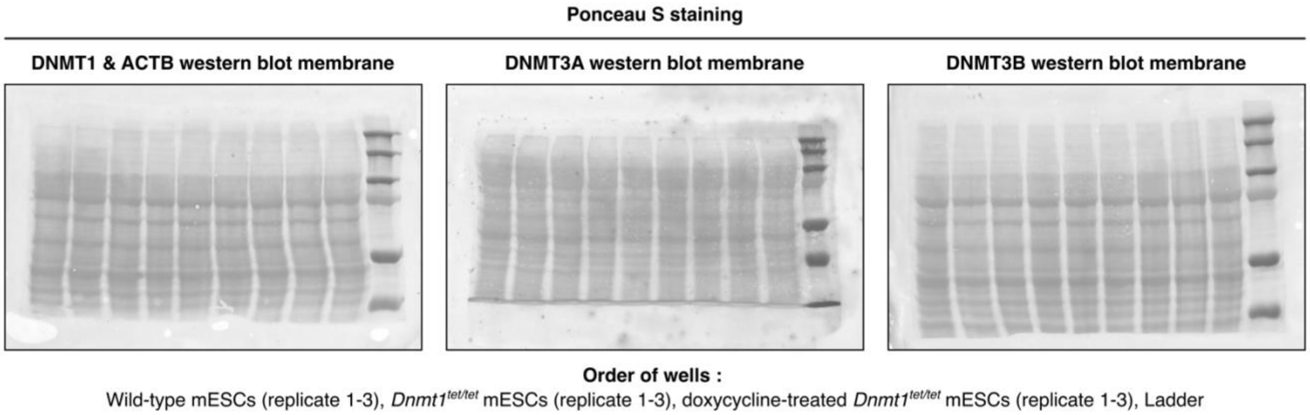


**Figure S1.** Perturbation of the DNA methylation machinery in *Dnmt1*$^{tet/tet}$ mESCs. Images of Ponceau S staining of western blot membranes used for normalizing the quantification of band intensities. *Related to Figure 1 and Data S2*.

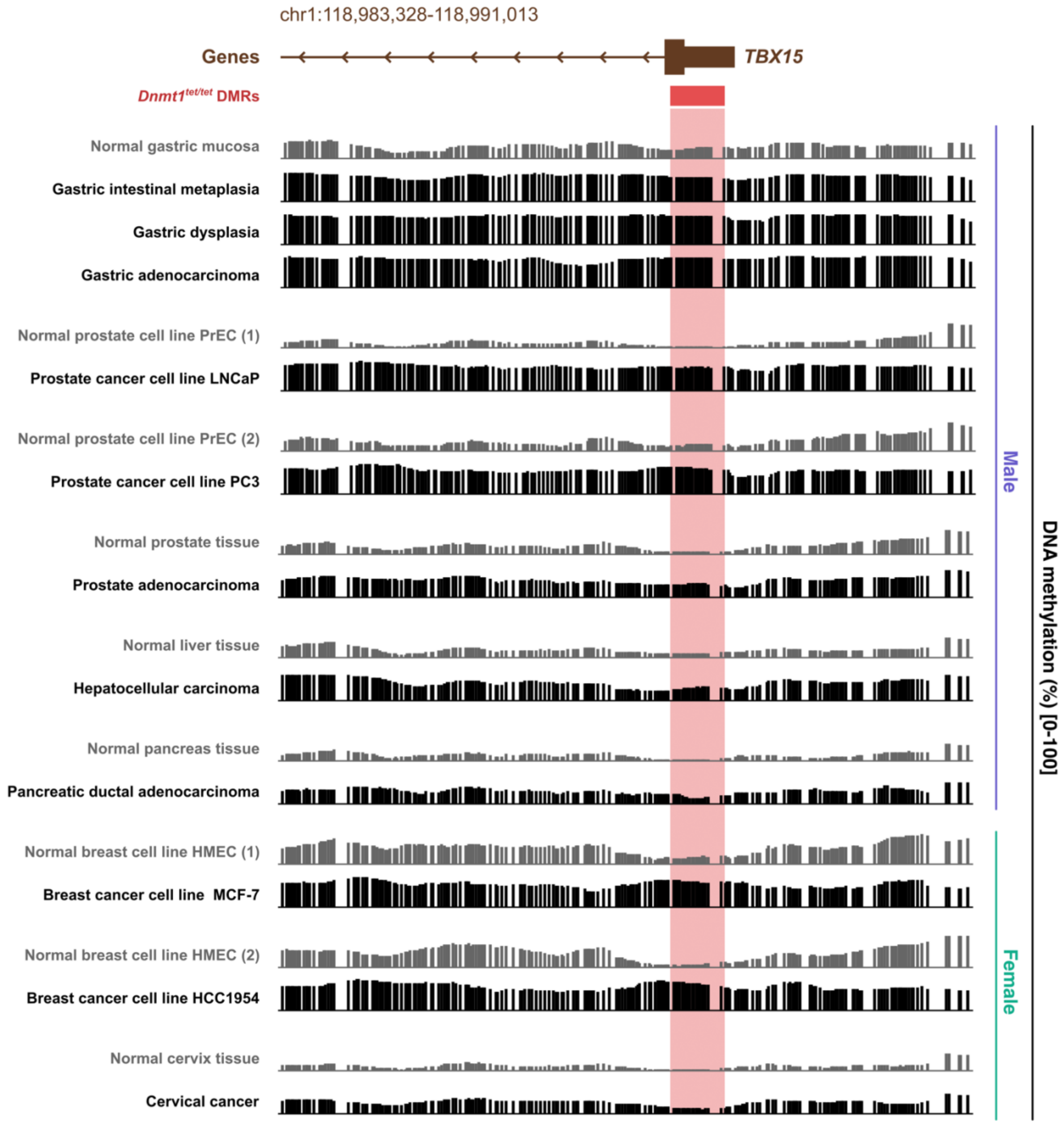


**Figure S2.** Conserved promoter hypermethylation between *Dnmt1*$^{tet/tet}$ mESCs and human cancer. Genomic tracks showing conserved *Dnmt1*$^{tet/tet}$ hypermethylation of the *TBX15* promoter across many human cancers.

| Genomic annotation | *Dnmt1*$^{tet/tet}$ differentially methylated regions (DMRs) | | |
|---|---|---|---|
| | Hypermethylated | Hypomethylated | Persistent hypermethylation |
| Genome wide | 4,303 | 81,144 | 1,708 |
| Core promoters | 2,536 | 2,307 | 1,232 |
| Proximal promoters | 673 | 1,952 | 284 |
| Distal promoters | 38 | 1,095 | 16 |
| Gene bodies | 589 | 28,544 | 95 |
| Intergenic | 467 | 47,246 | 81 |
| CpG islands | 3,426 | 440 | 1,631 |
| CGI shores | 97 | 4,605 | 34 |
| CGI shelves | 43 | 2,268 | 3 |
| Open sea | 737 | 73,831 | 40 |
| Imprinted regions | 2 | 15 | 0 |
| Retrotransposons | 383 | 32,778 | 5 |

**Table S1.** Number of *Dnmt1*$^{tet/tet}$ DMRs by genomic annotation that were either hypermethylated or hypomethylated and that showed persistent hypermethylation. Persistent hypermethylation: hypermethylated DMRs showing a sustained increase in DNA methylation ≥ 10% in doxycycline-treated *Dnmt1*$^{tet/tet}$ mESCs. *Related to Figures 2 and 5*.

| Human cell/tissue type | Sex | *Dnmt1*$^{tet/tet}$ hypermethylated promoter DMRs | | |
|---|---|---|---|---|
| | | Mapped | Hypermethylated | Hypomethylated |
| Gastric intestinal metaplasia (GIM) | Male | 3,149 | 430 (14%) | 9 (0.3%) |
| Gastric dysplasia (GD) | | | 436 (14%) | 20 (0.6%) |
| Gastric adenocarcinoma (GAC) | | | 537 (17%) | 16 (0.5%) |
| Prostate cancer cell line LNCaP (LNCaP) | Male | 3,171 | 278 (9%) | 22 (0.7%) |
| Prostate cancer cell line PC3 (PC3) | Male | 3,128 | 648 (21%) | 631 (20%) |
| Prostate adenocarcinoma (PRAD) | Male | 3,171 | 151 (5%) | 8 (0.3%) |
| Hepatocellular carcinoma (HCC) | Male | 3,171 | 291 (9%) | 23 (0.7%) |
| Renal cell carcinoma (RCC) | Male | 3,171 | 131 (4%) | 2 (0.1%) |
| Pancreatic ductal adenocarcinoma (PDAC) | Male | 3,167 | 329 (10%) | 19 (0.6%) |
| Breast cancer cell line MCF-7 (MCF-7) | Female | 2,915 | 687 (24%) | 358 (12%) |
| Breast cancer cell line HCC1954 (HCC1954) | Female | 3,116 | 468 (15%) | 13 (0.4%) |
| Cervical cancer (CC) | Female | 3,141 | 157 (5%) | 8 (0.3%) |
| Hepatocellular carcinoma (HCC) | Female | 3,136 | 182 (6%) | 63 (2%) |
| Renal cell carcinoma (RCC) | Female | 3,141 | 36 (1%) | 8 (0.3%) |
| Glioblastoma (GBM) | Female | 3,140 | 169 (5%) | 6 (0.2%) |

**Table S2.** Number of human genomic regions corresponding to the *Dnmt1*$^{tet/tet}$ hypermethylated promoter DMRs that were mapped and that were either hypermethylated or hypomethylated in various male and female human cancer samples. *See also Data S7. Related to Figures 6 and 7.*

| *Sample* | GIM | GD | GAC | LNCaP | PC3 | PRAD | HCC (M) | RCC (M) | PDAC | MCF-7 | HCC1954 | CC | HCC (F) | RCC (F) | GBM |
|---|---|---|---|---|---|---|---|---|---|---|---|---|---|---|---|
| GIM | 406 | 381 | 398 | 137 | 368 | 96 | 168 | 95 | 232 | 244 | 270 | 122 | 73 | 15 | 97 |
| GD | 381 | 431 | 413 | 136 | 385 | 93 | 168 | 94 | 237 | 248 | 281 | 126 | 78 | 17 | 94 |
| GAC | 398 | 413 | 513 | 163 | 447 | 107 | 189 | 97 | 257 | 290 | 318 | 133 | 90 | 19 | 102 |
| LNCaP | 137 | 136 | 163 | 277 | 174 | 115 | 124 | 69 | 118 | 177 | 203 | 58 | 63 | 20 | 76 |
| PC3 | 368 | 385 | 447 | 174 | 633 | 112 | 200 | 103 | 252 | 303 | 328 | 131 | 101 | 23 | 113 |
| PRAD | 96 | 93 | 107 | 115 | 112 | 148 | 83 | 48 | 84 | 107 | 131 | 41 | 39 | 15 | 48 |
| HCC (M) | 168 | 168 | 189 | 124 | 200 | 83 | 288 | 80 | 132 | 154 | 181 | 66 | 91 | 19 | 80 |
| RCC (M) | 95 | 94 | 97 | 69 | 103 | 48 | 80 | 129 | 86 | 90 | 103 | 49 | 47 | 23 | 58 |
| PDAC | 232 | 237 | 257 | 118 | 252 | 84 | 132 | 86 | 325 | 182 | 209 | 98 | 64 | 18 | 83 |
| MCF-7 | 244 | 248 | 290 | 177 | 303 | 107 | 154 | 90 | 182 | 687 | 284 | 108 | 76 | 19 | 90 |
| HCC1954 | 270 | 281 | 318 | 203 | 328 | 131 | 181 | 103 | 209 | 284 | 468 | 123 | 87 | 22 | 99 |
| CC | 122 | 126 | 133 | 58 | 131 | 41 | 66 | 49 | 98 | 108 | 123 | 157 | 36 | 11 | 44 |
| HCC (F) | 73 | 78 | 90 | 63 | 101 | 39 | 91 | 47 | 64 | 76 | 87 | 36 | 182 | 14 | 43 |
| RCC (F) | 15 | 17 | 19 | 20 | 23 | 15 | 19 | 23 | 18 | 19 | 22 | 11 | 14 | 36 | 22 |
| GBM | 97 | 94 | 102 | 76 | 113 | 48 | 80 | 58 | 83 | 90 | 99 | 44 | 43 | 22 | 169 |

**Table S3.** Number of regions of conserved $Dnmt1^{tet/tet}$ promoter hypermethylation that were shared between different human cancer samples. Regions on chromosomes X and Y were not included in this analysis. M: male, F: female. *See also Table S2*. *Related to Figure 7*.
